# BDNF/TrkB signaling endosomes in axons coordinate CREB/mTOR activation and protein synthesis in the cell body to induce dendritic growth in cortical neurons

**DOI:** 10.1101/2020.08.22.262923

**Authors:** Guillermo Moya-Alvarado, Miguel V. Guerra, Chengbiao Wu, William C. Mobley, Eran Perlson, Francisca C Bronfman

## Abstract

Brain-derived neurotrophic factor (BDNF) and its receptors tyrosine kinase receptor B (TrkB) and the p75 neurotrophin receptor (p75) are the primary regulators of dendritic growth in the central nervous system (CNS). After being bound by BDNF, TrkB and p75 are endocytosed into endosomes and continue signaling within the cell soma, dendrites, and axons. We studied the functional role of BDNF axonal signaling in cortical neurons derived from different transgenic mice using compartmentalized cultures in microfluidic devices. We found that axonal BDNF increased dendritic growth from the neuronal cell body in a cAMP response element-binding protein (CREB)-dependent manner. These effects were dependent on axonal TrkB but not p75 activity. Dynein-dependent BDNF-TrkB-containing endosome transport was required for long-distance induction of dendritic growth. Axonal signaling endosomes increased CREB and mTOR kinase activity in the cell body, and this increase in the activity of both proteins was required for general protein translation and the expression of Arc, a plasticity-associated gene, indicating a role for BDNF-TrkB axonal signaling endosomes in coordinating the transcription and translation of genes whose products contribute to learning and memory regulation.

## INTRODUCTION

Somatodendritic growth during central nervous system (CNS) development is essential for the establishment of proper connections (Jan and Jan, 2010). In addition, the morphology of dendritic arbors is maintained and undergoes plastic changes through a process that requires activity-dependent transcription factors (TFs), protein translation, and extracellular neurotrophic factors (Sutton and Schuman, 2006, Wong and Ghosh, 2002, Yap et al., 2018).

Brain-derived neurotrophic factor (BDNF) is a well-known neurotrophin whose transcription is regulated by neuronal activity and is required for the maintenance of dendrites of adult neurons in the CNS. BDNF binds two receptors, tyrosine kinase receptor B (TrkB) and the p75 neurotrophin receptor (p75). The classical trophic, growth-promoting actions of BDNF largely involve activation of TrkB, as mutant mice with inducible deletion of TrkB receptors exhibit a significant reduction in the number of dendritic arbors and degeneration of cortical neurons (Xu et al., 2000, Horch and Katz, 2002, Cheung et al., 2007, Yan et al., 1997, Huang and Reichardt, 2003).

Dendritic arborization mediated by TrkB activation in neurons is triggered by signaling pathways such as the ERK1/2 and the cAMP response element-binding protein (CREB) pathways (Finkbeiner et al., 1997; Xing et al., 1998, Gonzalez et al 2019). CREB also mediates activity-dependent transcription by regulating early gene expression (Flavell and Greenberg, 2008; Yap and Greenberg, 2018). In addition, activation of the class 1 phosphoinositide 3-kinase (PI3K)/Akt/mammalian target of rapamycin (mTOR) pathway increases the translation of a subset of mRNAs, including those that encode proteins related to BDNF-mediated regulation of dendritic size and shape in the CNS, in an mTOR-dependent manner (Takei et al., 2004, Ravindran et al., 2019, Schratt et al., 2004, Leal et al., 2014, Kumar et al., 2005, Minichiello et al., 2002, Dijkhuizen and Ghosh, 2005).

A large amount of evidence has shown that a complex of neurotrophins and neurotrophin receptors continue signaling inside the cell in specialized organelles named signaling endosomes, where neurotrophin receptors encounter specific signaling adaptors (Grimes et al., 1996, Bronfman et al., 2014, Debaisieux et al., 2016). Several studies have shown that endocytosis of TrkB is required for survival, dendritic arborization, and cell migration (Zheng et al., 2008, Zhou et al., 2007), indicating that endosomal signaling has a role in the physiological responses to TrkB. Consistent with these observations, studies from our laboratory have shown that there is a functional relationship between BDNF/TrkB signaling and the early recycling pathway in the regulation of dendritic morphology and CREB-dependent transcription of plasticity genes (Lazo et al., 2013, Moya-Alvarado et al., 2018, Gonzalez-Gutierrez et al., 2020).

Long-distance communication between neurotrophins in the target tissue and cell bodies has been described in the peripheral nervous system (PNS) and requires microtubule-associated molecular motors, such as cytoplasmic dynein (Bronfman et al., 2014, Reck-Peterson et al., 2018). During this process, NGF/TrkA signaling endosomes originating from the synaptic terminal propagate distal neurotrophin signals to the nucleus to allow transcriptional regulation, inducing neuronal survival (Cosker and Segal, 2014, Scott-Solomon and Kuruvilla, 2018, Riccio et al., 1997). Although it has been shown that BDNF is retrogradely transported along axons of cortical neurons, whether central neurons respond to distal neurotrophic signals is a relevant and understudied question in the neurobiology field. Indeed, no studies have directly assessed whether axonal BDNF signaling endosomes induce long-distance effects (Zhao et al., 2014, Olenick et al., 2019, Zhou et al., 2012, Stuardo et al., 2020, Cohen et al., 2011). Here, we studied whether axonal BDNF/TrkB signaling has a dendritic growth-promoting effect in the cell bodies of cortical neurons. We demonstrated for the first time that dynein-mediated transport of BDNF/TrkB signaling endosomes can modulate neuronal morphology. We also explored the BDNF/TrkB downstream signaling mechanism that regulates this process and showed that axonal BDNF-TrkB signaling endosomes coordinate the phosphorylation of CREB and PI3K-mTOR pathway-related proteins in cell bodies to increase protein translation and dendritic branching.

## RESULTS

### Axonal BDNF signaling promotes dendritic branching in a TrkB-dependent manner

Using cortical neurons in compartmentalized cultures in microfluidic chambers, we evaluated whether axonal stimulation with BDNF promotes dendritic arborization in a TrkB activity-dependent manner. We used two different systems: rat cortical neurons incubated with the tyrosine kinase inhibitor K252a (Tapley et al., 1992) and cortical neurons derived from TrkB^F616A^ knock-in mice. This mutation in TrkB enables pharmacological control of TrkB kinase activity using small molecule inhibitors, including 1-naphthylmethyl 4-amino-1-tert-butyl-3-(p-methyl phenyl) pyrazolo[3,4-d] pyrimidine (1NM-PP1), which selectively, rapidly, and reversibly inhibits TrkB activity. In the absence of such inhibitors, TrkB is fully functional (Chen et al., 2005). We also used cortical neurons derived from p75 exon III knockout mice (p75 KO^exonIII^) (Lee et al., 1992) to assess the dependence of dendritic growth and morphology on p75 expression and activity.

To evaluate morphological changes in neurons, we transfected cortical neurons in compartmentalized cultures (DIV 6) with plasmids expressing EGFP and then performed MAP2 immunostaining after fixing the neurons (Figure 1A). To identify the neurons that projected their axons to the axonal compartment (AC) and to assess leakage of media from the AC into the cell body compartment (CB), we used fluorescently labeled cholera toxin B subunit (Ctb). Ctb is internalized and retrogradely transported along neuronal axons and accumulates in the Golgi apparatus in the cell body (Escudero et al., 2019) (Figure 1A and Figure S1A and B). Cells that showed Ctb accumulation in the Golgi apparatus but not those in which Ctb diffusely labeled the soma were analyzed (Figure S1B). Only neurons in which the Golgi was labeled with Ctb were used for quantification. Since BDNF is also released from dendrites (Matsuda et al., 2009), axonal BDNF was selectively stimulated by incubating somas in the CB compartment with TrkB-Fc chimera (Shelton et al., 1995). TrkB-Fc neutralized the activity of endogenous BDNF released by neurons; when TrkB-Fc was not added to the CB compartment, axonal BDNF induced a nonsignificant increase in the number of distal dendrites (Figure S1C).

**Figure 1.**
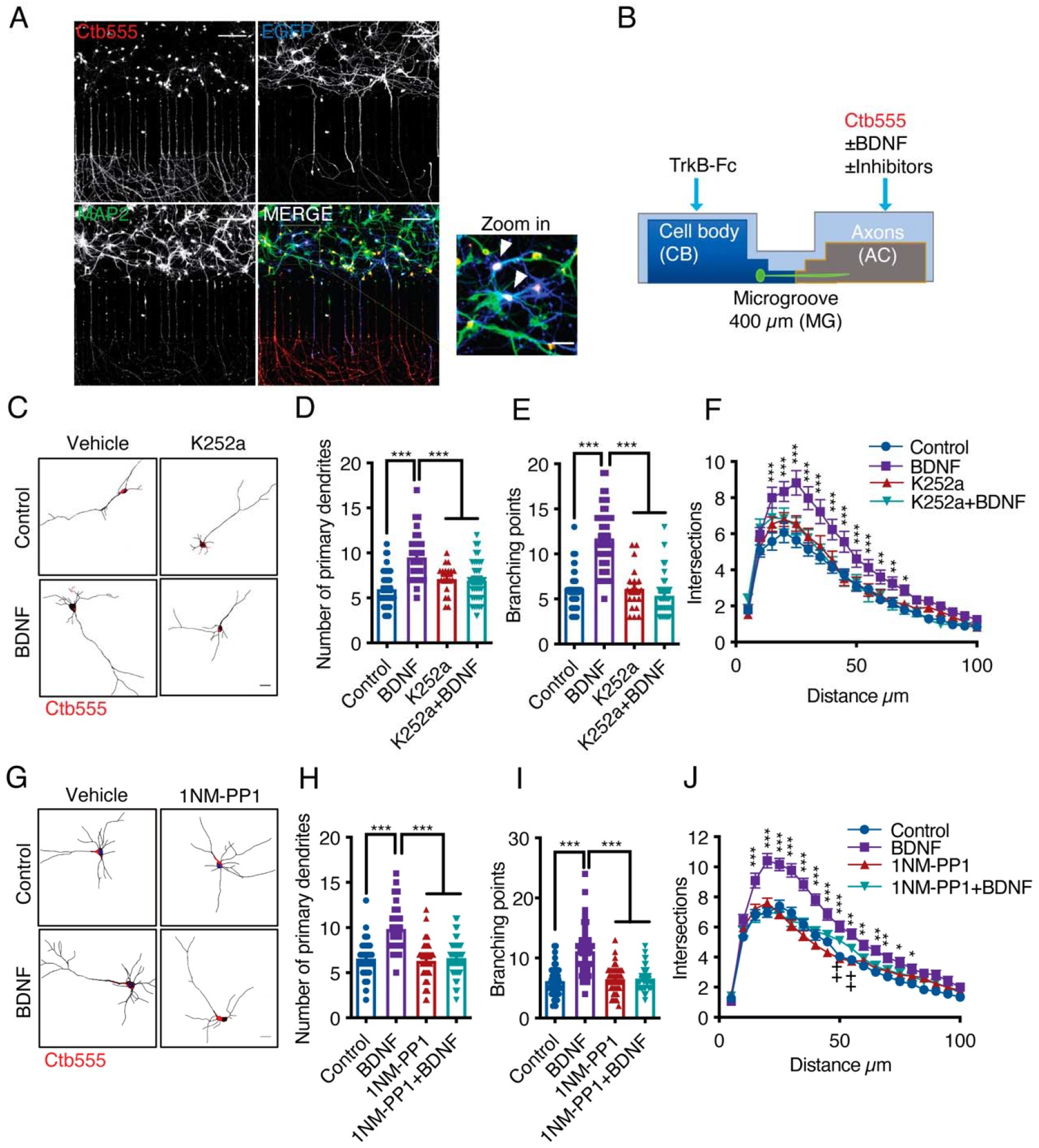
Addition of BDNF to axons promotes dendritic branching in a TrkB-dependent manner. **(A)** Representative images of compartmentalized microfluidic chambers (400 µm long and 15 µm wide microgrooves) after addition of BDNF/Ctb-555. Ctb-555 (upper left panel), EGFP (upper right panel), and MAP2 (lower left panel) were visualized by confocal microscopy (Scale bar = 50 µm) . The lower right panel is a merged image of the three signals. The white arrows in the zoomed-in panel indicate neurons costained with EGFP/MAP2/Ctb-555 (Scale bar = 20 µm), i.e., neurons that project axons through the microgrooves. **(B)** Experimental design used to study retrograde BDNF signaling. DIV six neurons were transfected with a plasmid expressing EGFP. TrkB-Fc (100 ng/mL) was applied to the CB. BDNF (50 ng/mL) in addition to fluorescently labeled Ctb (555) was added to the AC in the presence or absence of different inhibitors. BDNF and TrkB-Fc were administered for 48 hours to evaluate dendritic arborization. Finally, the neurons were fixed, and immunofluorescence for MAP2 was performed. **(C)** Representative images of the CB (red indicates Ctb-555) of compartmentalized rat cortical neurons whose axons were treated with DMSO (control), K252a (200 nM), BDNF (50 ng/mL) or BDNF following preincubation with K252a **(D-F).** Quantification of primary dendrites **(D)** and branching points **(E)** and Sholl analysis **(F)** for neurons labeled with EGFP/MAP2/Ctb-555 under each experimental condition. n= 40-45 neurons from 3 independent experiments. **(G)** Representative images of the CB (red indicates Ctb-555) of compartmentalized mouse TrkB^F616A^ cortical neurons whose axons were treated with DMSO (control), 1NM-PP1 (1 µm), BDNF (50 ng/mL), or BDNF following preincubation with 1NM-PP1. **(H-J)** Quantification of primary dendrites **(H)** and branching points **(I)** and Sholl analysis **(J)** for neurons labeled with EGFP/MAP2/Ctb-555 under the four different experimental conditions described in G. Scale bar = 20 µm. n= 40-45 neurons from 3 independent experiments. *p<0.05, **p< 0.01, ***p< 0.001, ++p<0.01, the 1NM-PP1 group vs. the 1NM-PP1+BDNF group. The data were analyzed by one-way ANOVA followed by Bonferroni’s multiple comparisons post hoc test (D, E, H and I). Two-way ANOVA followed by Bonferroni’s multiple comparisons post hoc test (F and J). The results are expressed as the mean ± SEM.

The experiment presented in Figure 1B and explained above revealed that the addition of BDNF to axons led to increases in the numbers of primary dendrites, branching points and overall dendritic arbors in the cell bodies of cortical neurons from rats (Figure 1C-F) and mice (Figure 1G-J). Furthermore, we found that this effect required axonal TrkB activation, as treating the axons of rat cortical neurons with K252A or treating cortical neurons from TrkB^F616A^ knock-in mice with 1NM-PP1 abolished the effect of axonal BDNF on neuronal morphology (Figure 1C-J).

To evaluate the contribution of p75 to the dendritic arborization induced by axonal BDNF, we used cortical neurons from p75 KO^exonIII^ mice. Using the same experimental design described in Figure 1A and B, we prepared cultures derived from p75 KO^exonIII^ (p75^+/+^), heterozygous (p75^+/-^), and homozygous p75 KO^exonIII^ (p75^-/-^) mice. We observed that the number of branching points and primary dendrites induced by axonal stimulation with BDNF was unchanged by the p75 genotype (Figure 2A-C). Nevertheless, Sholl analysis of dendritic morphology revealed that the basal morphology of the dendritic arbors was altered in the absence of p75. While p75^-/-^ neurons possessed a smaller number of primary dendrites, they had an increased number of distal dendrites (Figure 2D); however, this phenotype was rescued by axonal BDNF (Figure 2E). Altogether, these results indicate that stimulation of the axons of cortical neurons with BDNF increases dendritic branching and that this effect depends mainly on the activity of TrkB in axons.

**Figure 2.**
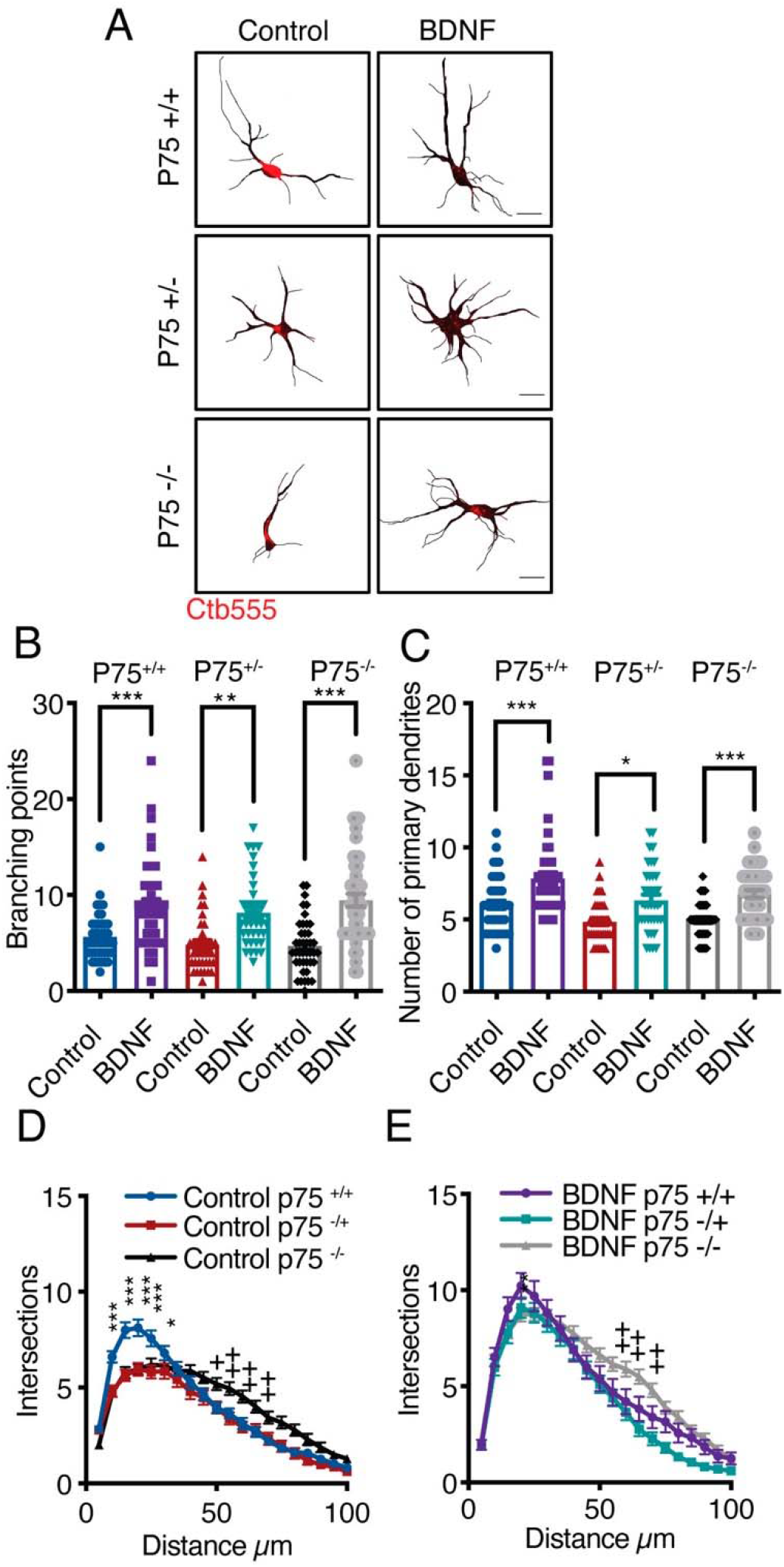
Axonal BDNF increases dendritic arborization in a p75-independent manner. **(A)** Representative images of the CB (red indicates Ctb-555) of compartmentalized p75^+/+^, p75^+/-^ or p75^-/-^ mouse cortical neurons whose axons were treated with BDNF (50 ng/mL). Quantification of branching points **(B)** and primary dendrites **(C)** in MAP2/Ctb-555-labeled neurons from mice of the different genotypes described in A. **(D)** Sholl analysis of cultured neurons from mice of each genotype following application of TrkB-Fc to the CB (nonstimulated). **(E)** Sholl analysis of cultured neurons from mice of each genotype following addition of TrkB-Fc to the CB and BDNF to the AC. n= 40-50 neurons from 2 independent experiments (3 mice per experiment). Scale bar = 10 µm *p<0.05, **p< 0.01, ***p< 0.001, ++p<0.01, neurons from p75^-/-^ mice vs. neurons from p75^+/+^ mice. The data were analyzed by one-way ANOVA followed by Bonferroni’s multiple comparisons post hoc test (B and C). Two-way ANOVA followed by Bonferroni’s multiple comparisons post hoc test (D and E). The results are expressed as the mean ± SEM.

### CREB activity is required for long-distance BDNF signaling

Several groups have shown that axonal BDNF signaling promotes nuclear CREB phosphorylation in different neuronal models (Zhou et al., 2012, Watson et al., 1999, Deinhardt et al., 2006, Gonzalez-Gutierrez et al., 2020); however, the physiological relevance of CREB activation in this process has not been evaluated. To evaluate CREB activation, we used a well-characterized polyclonal antibody against CREB phosphorylated at S133 (Lonze and Ginty, 2002, Gonzalez-Gutierrez et al., 2020) and studied the time course of CREB phosphorylation induced by axonal BDNF treatment. At DIV 5, we incubated the AC with Ctb-555 overnight to identify neurons with axons in the AC (Figure 3A). BDNF induced an increase in nuclear CREB phosphorylation as soon as 30 min, with nuclear CREB phosphorylation peaking at two time points: at 30 min and 180 min (Figure 3A and B).

**Figure 3.**
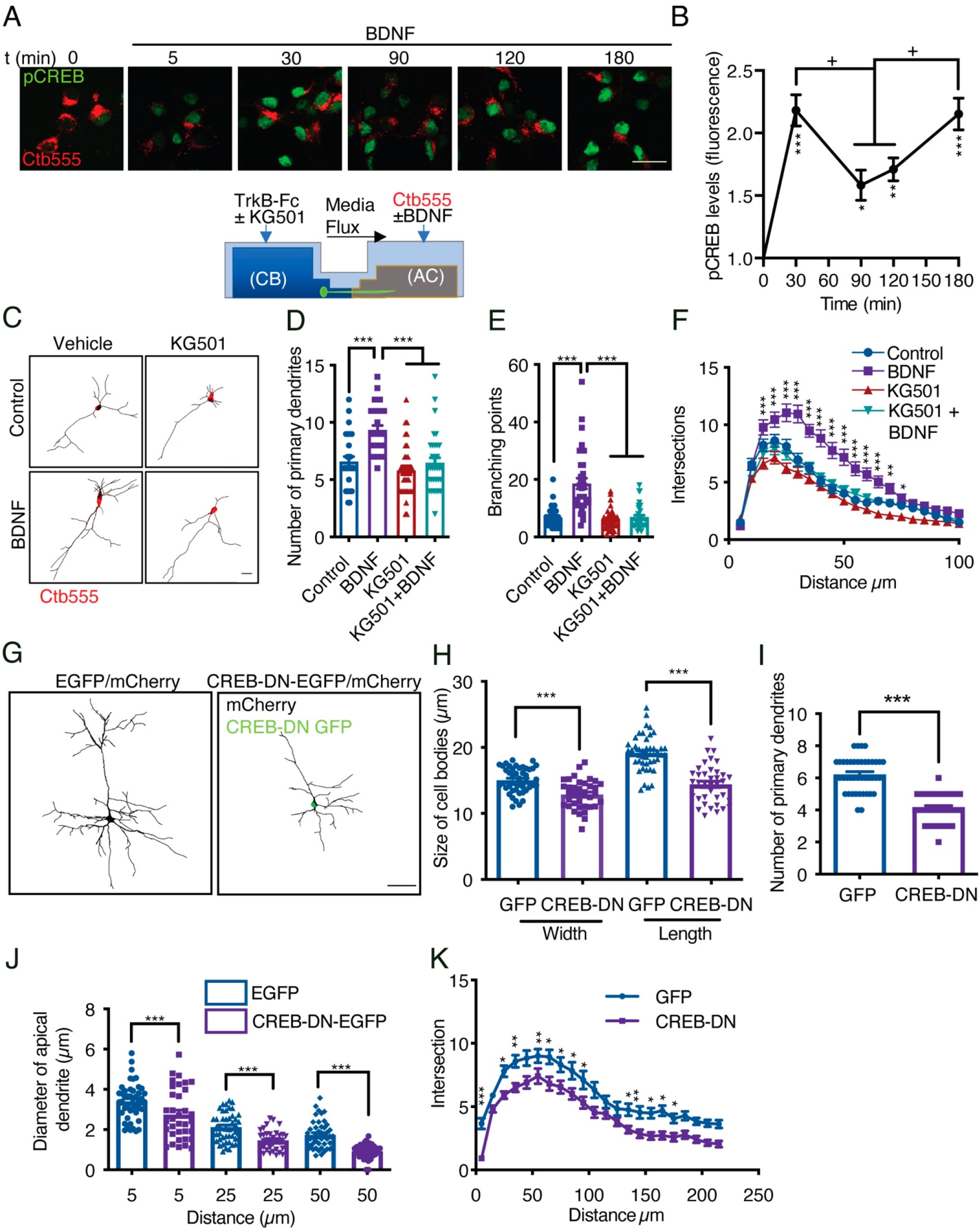
CREB activity is required for long-distance axonal BDNF-induced dendritic arborization. **(A)** Lower panel, schematic representation of the protocol used to evaluate CREB phosphorylation induced by axonal BDNF and the requirement of CREB for dendritic arborization induced by BDNF. Upper panel, representative figures of nuclear activated CREB (pCREB, S133) immunostaining in control neurons and neurons whose axons were stimulated with BDNF for different times. Ctb-555 was added to the AC of compartmentalized neurons overnight, and after 90 min of serum deprivation, BDNF (50 ng/mL) was added to the AC for 5, 30, 90, 120 or 180 min. pCREB is shown in green and Ctb-555 is shown in red. Scale bar = 10 µm. **(B)** Quantification (arbitrary units, A.U.) of pCREB fluorescence in the nucleus in the presence and absence of BDNF. Only neurons positive for Ctb-555 were included in the analysis. n= 30-40 neurons from 2 independent experiments. *p<0.05, **p< 0.01, ***p< 0.01 vs. the control group; + p<0.05 vs. the 90-min and 120-min BDNF treatment groups. The results are expressed as the mean ± SEM. The data were analyzed by one-way ANOVA followed by Bonferroni’s multiple comparisons post hoc test. **(C)** Representative images of the CB (red indicates Ctb-555) of compartmentalized neurons whose cell bodies were treated with DMSO (control), KG501, BDNF or BDNF in the presence of KG501. TrkB-Fc (100 ng/mL) and KG501 (10 µM) were applied to the CB for 1 hour. Then, BDNF (50 ng/mL) was added to the AC for 48 hours in the presence of Ctb-555, and the CB was treated with or without KG501. Finally, the neurons were fixed, and immunofluorescence for MAP2 was performed. Scale bar = 20 µm. **(D-F)** Quantification of primary dendrites **(D)** and branching points **(E)** and Sholl analysis **(F)** for neurons labeled with EGFP/MAP2/Ctb-555 under each experimental condition. n= 30-35 neurons from 3 independent experiments. *p<0.05, **p< 0.01, ***p< 0.001. The data were analyzed by one-way ANOVA followed by Bonferroni’s multiple comparisons post hoc test (D and E). Two-way ANOVA followed by Bonferroni’s multiple comparisons post hoc test was used to analyze the Sholl analysis data (F). The results are expressed as the mean ± SEM. **(G)** Representative images of pyramidal neurons from cortical layers II-III of the sensory motor cortex cotransduced with AAV1 expressing CREB-DN-EGFP or EGFP and mCherry. mCherry was amplified by immunostaining for the fluorescent protein. Scale bar = 50 µm. Neurons expressing CREB-DN-EGFP showed reductions in size **(H),** the number of primary dendrites **(I)**, the apical dendrite diameter **(J)**, and branching, as measured by Sholl analysis **(K)**. The results are expressed as the mean ± SEM. The data were analyzed by one-way ANOVA followed by Bonferroni’s multiple comparisons post hoc test (H-J). Two-way ANOVA followed by Bonferroni’s multiple comparisons post hoc test was used to analyze the Sholl analysis data (K). n=35-41 neurons per experimental group derived from 5 different mice.

To confirm that CREB activity is required for dendritic arborization, we stimulated the axons of cortical neurons whose cell bodies had been incubated with KG501 for 48 hours with BDNF (Figure 3C). KG501 is a small molecule that disrupts the interaction between CREB and CREB-binding protein (CBP) (Best et al., 2004). We found that KG501 abolished the increase in dendritic arborization induced by treating axons with BDNF (Figure 3D-G). We also evaluated whether p75 is required for the phosphorylation of CREB upon BDNF stimulation. Consistent with the results presented in Figure 2, the absence of p75 did not impact BDNF-induced CREB phosphorylation (Figure S2A and B), indicating that in our model, p75 was not required for BDNF-induced CREB phosphorylation.

Cortical neurons in the cortico-callosal pathway possess long axons (Figure S2 C), and this pathway is known to be modulated by BDNF and TrkB (Andreska et al., 2020, Shimada et al., 1998). When cortical neurons were transduced with adeno-associated virus (AAV)1 expressing dominant negative CREB (CREB-DN) for 3 weeks, the cell body size, the number of neurons, the apical dendrite diameter and branching were reduced (Figure 3H-L), and nonapoptotic cell death, as indicated by fragmentation of the nucleus revealed by Hoechst staining, was observed (Figure S2D), indicating that CREB is required for the maintenance of neuronal morphology *in vivo* but had no effects on neuronal survival at the time point studied.

Since CREB has been widely described as a significant regulator of neurotrophin-induced survival of PNS neurons (Finkbeiner et al., 1997, Bonni et al., 1999, Lonze and Ginty, 2002), we assessed whether KG501 has an effect on neuronal survival under our experimental conditions by assessing neuronal apoptosis using TUNEL staining. We found that after 48 hours of treatment, KG501 did not affect neuronal survival. We used oligomycin A, an inhibitor of oxidative phosphorylation, as a positive control to induce apoptotic death of cortical neurons (Figure S2 E and F).

Our results suggest that axonal BDNF promotes CREB phosphorylation, and that CREB-dependent transcription is required for long-distance induction of dendritic branching by BDNF.

### Dynein-dependent transport of endocytosed TrkB in axons is required for long-distance BDNF signaling

In axons, long-range retrograde transport of organelles and signaling molecules is achieved by the minus-end microtubule molecular motor cytoplasmic dynein (Hirokawa et al., 2010). Several groups have shown that in CNS neurons, BDNF and TrkB are efficiently retrogradely transported (Zhou et al., 2012, Olenick et al., 2019). To further confirm that this process is dynein dependent, we monobiotinylated BDNF *in vitro* and labeled it with streptavidin quantum dots (QDs) for *in vivo* imaging (Stuardo et al., 2020). We found that BDNF-QDs was efficiently targeted to the retrograde transport pathway and that the retrograde transport of BDNF-QDs was reduced by approximately 80% by inhibition of dynein with ciliobrevin D (CilioD), a specific inhibitor of dynein ATP activity (Sainath and Gallo, 2015) (Figure 4A and B). To evaluate whether axonal dynein inhibition reduces the transport of axonal TrkB, we evaluated axonal BDNF-induced nuclear phosphorylation of CREB and dendritic arborization in axons in the presence and absence of CilioD (Figure 4C). We found that CilioD reduced CREB phosphorylation (Figure 4D and E) and dendritic arborization induced by the application of BDNF to axons (Figure 4G-J), which is consistent with the concept that CREB-dependent arborization induced by axonal BDNF is dependent on dynein transport.

**Figure 4.**
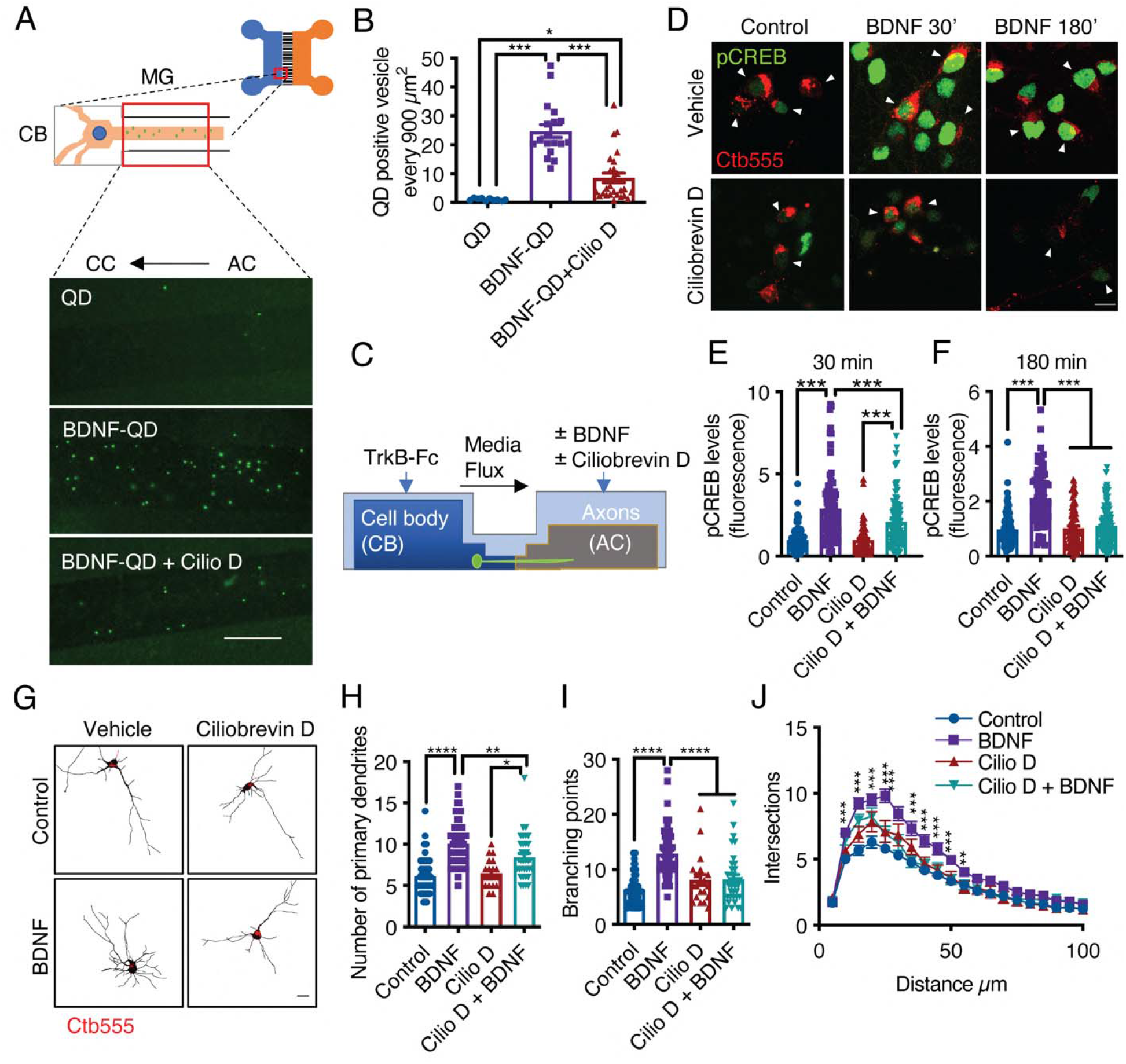
Dynein activity is required for phosphorylation of CREB and dendritic arborization induced by BDNF. **(A)** Schematic representation of the protocol used to evaluate the retrograde transport of BDNF. Biotinylated BDNF conjugated to streptavidin QD-655 (2 nM) was added to AC for 4 hours in the presence or absence of CilioD (20 µM). Then, after washing, the neurons were fixed, and the accumulation of BDNF-QDs in the proximal compartment of the microgrooves (MG) was evaluated. Unconjugated QDs were used as controls. Representative images of BDNF-QDs (green) in the proximal region of the MG under each condition. Scale bar = 10 µm **(B)** Quantification of QD-positive vesicles in every 900 µm^2^ area of the proximal region of MG under each condition. N= 36 MG from 3 independent experiments were analyzed. **(C)** Experimental design used to study dynein-dependent axonal BDNF signaling in compartmentalized cortical neurons. DIV five cortical neurons were retrogradely labeled with Ctb-555 (red) overnight. The next day, the culture medium was replaced with serum-free medium for 90 min, and CilioD was applied to the AC. Then, BDNF (50 ng/mL) was added to the AC for 30 or 180 min in the presence or absence of CilioD. Finally, the cultures were fixed, and pCREB (S133) was visualized by immunofluorescence. **(D)** Representative images of nuclear pCREB in neurons. Scale bar = 10 µm. **(E-F)** Quantification of (arbitrary units, A.U.) pCREB immunostaining in the nuclei of neurons labeled with Ctb-555 and stimulated with BDNF for 30 min (E) or 180 min (F). n= 83-111 neurons from 3 independent experiments. **(G)** DIV six cortical neurons were transfected with EGFP, TrkB-Fc (100 ng/mL) was applied to the CB, and Ctb-555 and BDNF (50 ng/mL) with or without CilioD (20 µM) were added to the AC for 48 hours. Finally, the neurons were fixed, and immunofluorescence for MAP2 was performed. Representative images of the morphology of compartmentalized neurons that were labeled with Ctb-555 (red) and whose axons were treated with DMSO (control), CilioD, BDNF or BDNF following pretreatment with CilioD for 48 hours. Scale bar = 20 µm. **(H-J)** Quantification of primary dendrites **(H)** and branching points **(I)** and Sholl analysis **(J)** for neurons labeled with EGFP/MAP2/Ctb-555 under each experimental condition described in G. n= 34-65 neurons from 3 independent experiments. *p<0.05, **p< 0.01, ***p< 0.001, ****p<0.0001. The results are expressed as the mean ± SEM. The data were analyzed by one-way ANOVA followed by Bonferroni’s multiple comparisons post hoc test (E, F, H, and I). The Sholl analysis data were analyzed by two-way ANOVA followed by Bonferroni’s multiple comparisons post hoc test.

### The somatodendritic activity of axonal TrkB is required for long-distance signaling triggered by BDNF in axons

Several reports have shown that endocytosed axonal BDNF is retrogradely transported to and accumulates in cell bodies (Olenick et al., 2019; Zhou et al., 2012; Xie et al, 2012). These findings, together with the observations presented in this research, suggest that BDNF is transported along with TrkB via signaling endosomes to the cell body, where the BDNF-TrkB complex continues to signal.

To our knowledge, there is no evidence that active TrkB in axons, reaches the cell body and continues to signal to induce CREB activation. Thus, we design an experiment where we ought to activate TrkB in TrkB^F616A^ knock-in mouse neurons in compartmentalized cultures by applying BDNF to the AC and inhibited the activity of TrkB upon its arrival in the cell body by applying 1NM-PP1 to the CB. To achieve this goal, we first studied whether NM-PP1 can reduce TrkB activity after BDNF has activated the receptor in noncompartmentalized cultures. We stimulated neurons with BDNF for 30 min and then incubated them with 1NM-PP1 or DMSO for an additional 30 min. As a control, we preincubated the neurons with 1NM-PP1 for 1 hour to prevent the activation of TrkB (Figure S3A). As expected, preincubation of neurons with 1NM-PP1 prevented the activation of TrkB and Akt induced by BDNF (Figure S3B). In addition, we observed that treatment with 1NM-PP1 after BDNF stimulation decreased the phosphorylation of TrkB and Akt to levels similar to those before incubation (Figure S3B-D). Consistently, immunofluorescence revealed that 1NM-PP1 treatment after BDNF stimulation reduced the nuclear phosphorylation of CREB and the phosphorylation of 4E-BP1 and S6 ribosomal protein (S6r), two proteins downstream of mTOR activation induced by BDNF (Figure S3E-H)). These results indicate that 1NM-PP1 can turn off TrkB after it is activated by BDNF.

Then, to evaluate whether axonal-derived activated TrkB reaches the cell body and is required for the propagation of axonal BDNF signaling, we performed transfection, pulse chase and NM-PP1 inhibition experiments as follows. We first transfected neurons with a plasmid expressing TrkB containing a Flag NH2-terminal epitope (Flag-TrkB) (Lazo et al., 2013) and then incubated the axons with a Flag-specific antibody at 4°C, washed them, and treated them with BDNF at 37°C (Figure 5A). After fixation, we evaluated the colocalization of Flag with pTrkB in microgrooves in proximity to the AC (distal microgrooves), in the vicinity of the CB (proximal microgrooves) and in the CB (Figure 5B). We found that signaling endosomes containing activated TrkB move along axons and reach the cell body upon stimulation of axons with BDNF. Subsequently, we directly assessed whether we can reduce TrkB activity in the cell bodies of TrkB^F616A^ mouse-derived neurons after stimulation of axons with BDNF. Somatodendritic TrkB activity was evaluated after applying 1NM-PP1 to the CB or AC (Figure 5C). Application of BDNF to the axons increased the immunostaining intensity of endogenous pTrkB in the CB (Figure 5D and E). Consistent with the observations in noncompartmentalized neuronal cultures, the application of 1NM-PP1 to the AC completely abolished the accumulation of pTrkB in the cell body (Figure 5D and E). Furthermore, the application of 1NM-PP1 to the cell body decreased the amount of activated TrkB in the cell body after stimulation of axons with BDNF (Figure 5D and E).

**Figure 5.**
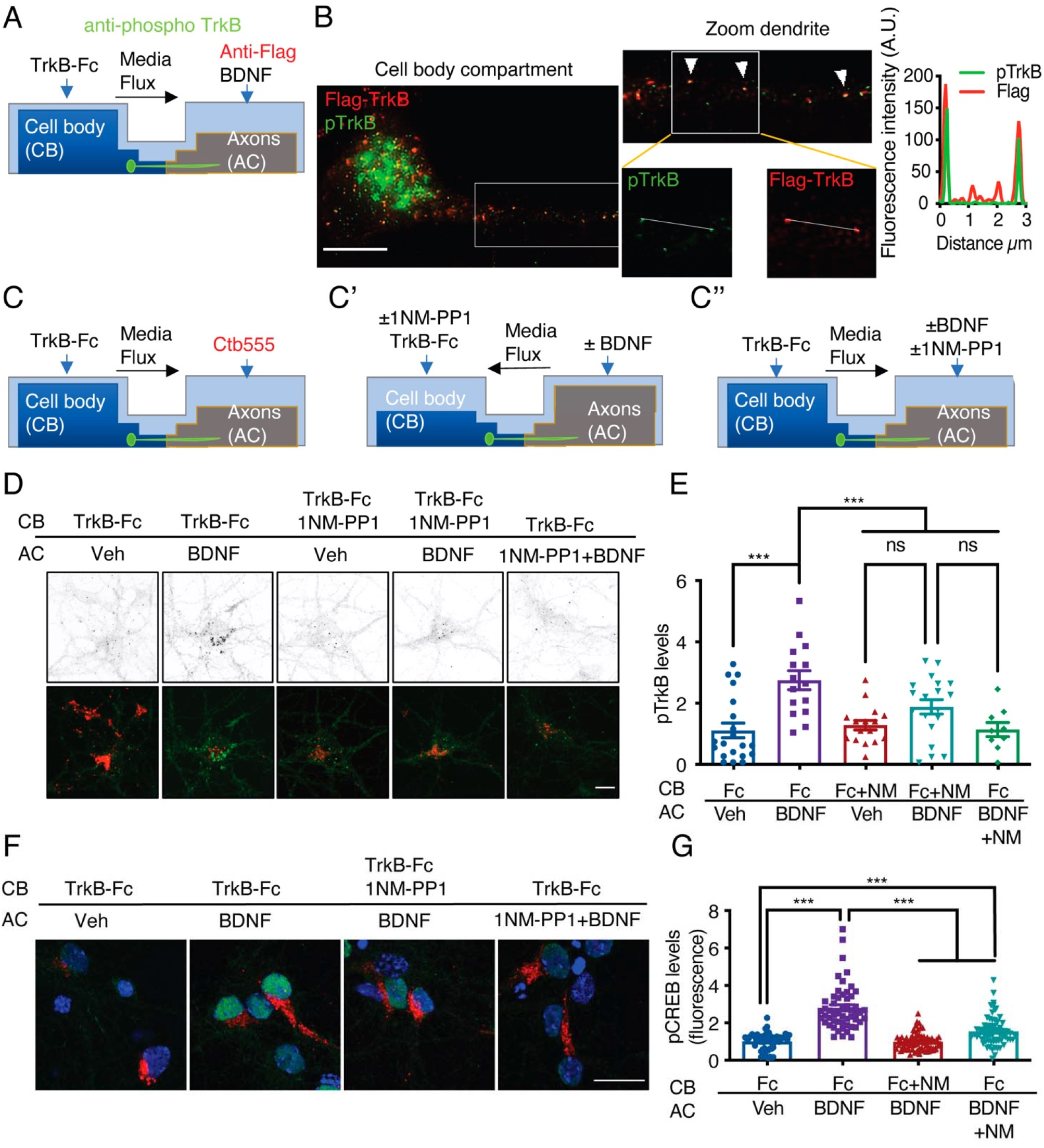
Somatodendritic activity of axonal TrkB is required for long-distance BDNF signaling. **(A)** Schematic representation of the protocol used to label axonal TrkB. DIV five compartmentalized cortical neurons were transfected with Flag-TrkB for 48 hours. At DIV seven, anti-Flag antibody was added to AC for 40 min at 4°C. Then, neurons were incubated with or without BDNF (50 ng/mL) at 37°C. Finally, immunofluorescence for pTrkB (Y816, pTrkB) (green) and anti-Flag (red) was performed. **(B)** Immunofluorescence of pTrkB and internalized Flag-TrkB in compartmentalized neurons whose axons were treated with BDNF, as shown in A. Right panel, representative images of neurons whose axons were treated with BDNF. Left panel, a magnified image of a proximal neuronal dendrite. The arrows indicate colocalization of Flag (red) and pTrkB (green). Scale bar = 5 μm. Right panel, graphs showing the fluorescence intensity pixel by pixel along the white lines shown in the right panels in B. The green line indicates pTrkB fluorescence, and the red lines indicate Flag-TrkB fluorescence. **(C-C’’)** Schematic representation of the protocol used to stimulate neurons in D-G. **(C)** At DIV 6, Ctb-555 was added to the AC of cortical neurons from TrkBF616A mice overnight. The next day (DIV 7), the neurons were subjected to two treatments: (C’) depletion of B27 supplement for 1 hour in the presence of 1NM-PP1 (1 μM) in the CB, with flux toward the CB. Then, TrkB-Fc (100 ng/mL) and 1NM-PP1 were added to the CB, and BDNF (50 ng/mL) was applied or not to the AC for 3 hours. (C’’) Neurons were depleted of B27 supplement for 1 hour in the presence or absence of 1NM-PP1 (1 μM) in the AC, with flux toward the AC. Then, TrkB-Fc was applied to the CB, and BDNF was added or not to the AC in the presence or absence of 1NM-PP1 for 3 hours. Finally, the neurons were fixed, and immunofluorescence for pTrkB (Y816) or pCREB (S133) was performed. **(D)** Representative images of pTrkB in the CB compartment of Ctb-positive neurons treated as described in A. Scale bar = 5 µm. Quantification of pTrkB levels in the cell body in each treatment group described in B. n=3 independent experiments. **(F)** Representative image of pCREB immunostaining in cortical neurons whose axons were stimulated with BDNF for 3 hours in the presence or absence of 1NM-PP1 in the CB (1NM-PP1/BDNF) or AC (1NM-PP1 + 1NM-PP1). Scale bar = 10 µm. **(G)** Quantification of pCREB levels in Ctb-positive neurons under each condition. n= 78-86 neurons from 3 independent experiments. **p< 0.01, ***p<0.001. vs. the control group. The results are expressed as the mean ± SEM. The data were analyzed by one-way ANOVA followed by Bonferroni’s multiple comparisons post hoc test.

These results suggest that 1NM-PP1 inhibited the phosphorylation of endosomal TrkB after it arrived in the neuronal cell body. To further confirm this phenomenon, we studied the phosphorylation of CREB in the cell bodies of TrkB^F616A^ mouse neurons in compartmentalized cultures after stimulation of axons with BDNF in the presence or absence of 1NM-PP1. We found that somatodendritic TrkB activity was required for CREB phosphorylation induced by stimulation of axons with BDNF (Figure 5F and G). Together, our findings indicate that signaling endosomes carrying activated TrkB are required for the propagation of axonal BDNF signals to the nuclei of CNS neurons.

### The somatodendritic PI3K/Akt/mTOR pathway is required for the induction of dendritic branching by axonal BDNF

BDNF-mediated dendritic arborization is achieved via activation of the PI3K/Akt/mTOR signaling pathway, which increases protein translation in the cell body and dendrites (Dijkhuizen and Ghosh, 2005, Kumar et al., 2005). Additionally, PI3K has been shown to mediate retrograde NGF signaling in sympathetic neurons (Kuruvilla et al., 2000). Furthermore, it was reported that axonal endosomes are positive for TrkB and Akt (Goto-Silva et al., 2019), suggesting that axonal PI3K/Akt signaling may contribute to long-distance dendritic arborization induced by BDNF. Thus, we applied BDNF to the the AC of compartmentalized cortical neurons in the presence or absence of LY294002, a potent general PI3K inhibitor (Figure 6A). Interestingly, the application of LY294002 to axons did not affect dendritic arborization induced by axonal BDNF (Figure 6B-E). Confirming previous results (Finsterwald et al; 2010), LY294002 reduced BDNF-induced dendritic arborization in noncompartmentalized cortical cultures, indicating that LY294002 exerted an effect in cortical neurons (Figure S4A and B). Furthermore, inhibition of PI3K activity in axons did not reduce the transport of BDNF-QDs (Figure 6F and G). This result indicates that PI3K activity is not required for the retrograde transport of axonal BDNF signals. Therefore, we investigated whether PI3K signaling in the somatodendritic compartment is required for axonal BDNF-induced dendritic arborization by applying LY294002 to the CB of compartmentalized cortical cultures (Figure 6H). We observed that the application of LY294002 to the CB inhibited dendritic arborization induced by axonal BDNF (Figure 6I-L), suggesting that PI3K activation in the cell body is required for long-distance signaling induced by axonal BDNF.

**Figure 6.**
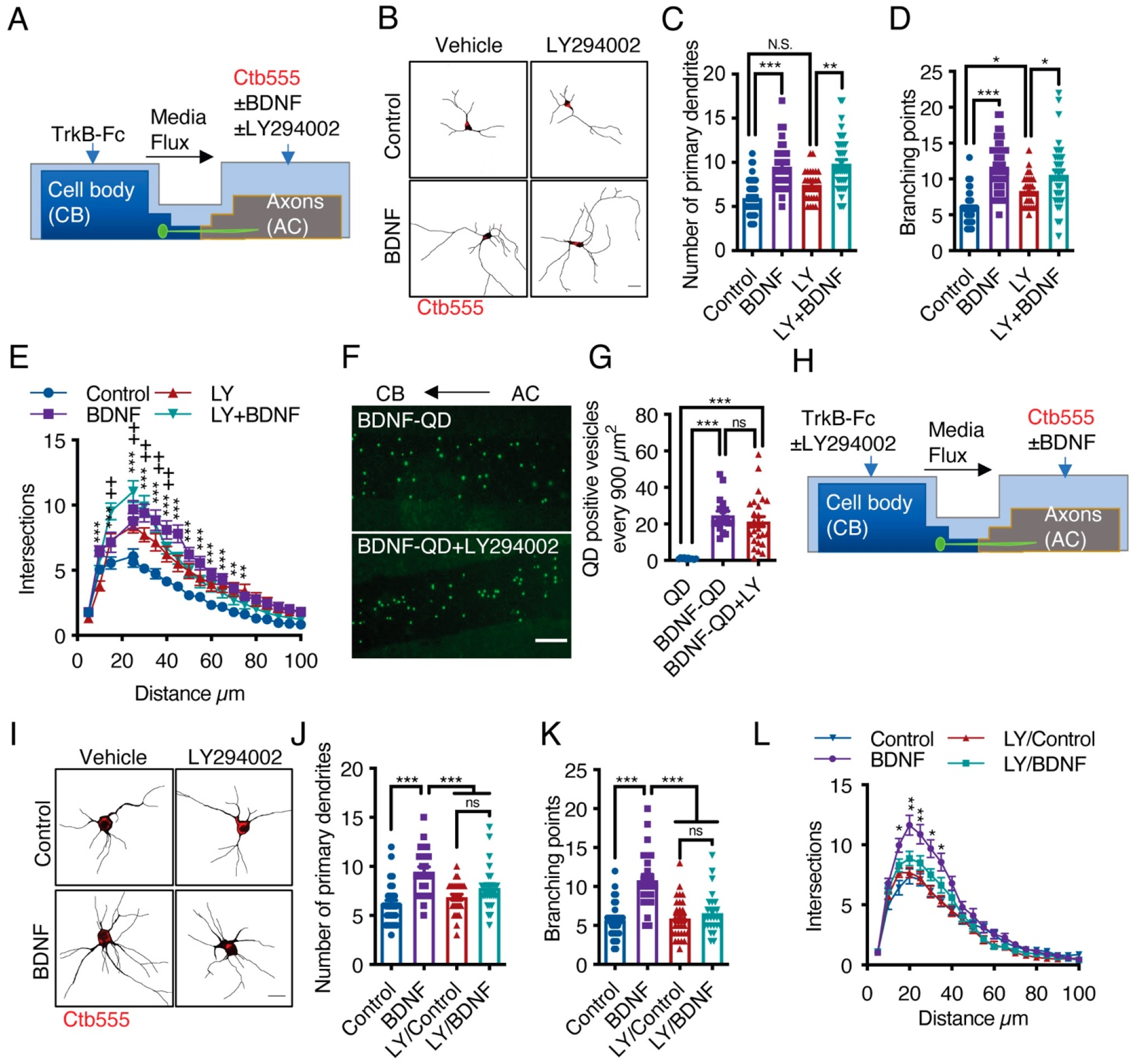
Activation of PI3K activity in the cell body but not the axons is required for dendritic arborization induced by axonal BDNF. **(A)** Schematic representation of the protocol used to stimulate compartmentalized cortical neurons. DIV six cortical neurons were transfected with EGFP, and then TrkB-Fc (100 ng/mL) was added to the CB and Ctb-555 and BDNF (50 ng/mL) with or without LY294002 (10 µM) were applied to the AC for 48 hours. Finally, the neurons were fixed, and immunofluorescence for MAP2 was performed. **(B)** Representative images of the CB (red indicates Ctb-555) of compartmentalized neurons whose axons were treated with DMSO in the presence or absence of BDNF and LY294002. Scale bar 20 µm. **(C-E)** Quantification of primary dendrites **(C)** and branching points **(D)** and Sholl analysis **(E)** for neurons labeled with EGFP/MAP2/Ctb-555 under the different experimental conditions described in B. n= 29-48 neurons from 3 independent experiments. **(F)** BDNF-QDs were added to the AC for 4 hours in the presence or absence of LY294002 (10 µM) to promote the retrograde transport of BDNF. Then, the neurons were fixed, and the accumulation of BDNF-QDs in the proximal compartment of the microgrooves (MG) was evaluated. Representative image of accumulated BDNF-QDs (green dots) in the proximal part of MG under each treatment. Scale bar = 10 µm. **(G)** Quantification of QD-positive vesicles in every 900 µm^2^ area of the proximal region of MG under each condition. Scale bar = 10 µm. n=36 MG from 3 independent experiments were analyzed. **(H)** Schematic representation of the protocol used to stimulate compartmentalized cortical neurons. DIV six cortical neurons were transfected with EGFP. Then, TrkB-Fc (100 ng/mL) was added to the CB in the presence or absence of LY294002 (10 µM), and Ctb-555 and BDNF (50 ng/mL) were applied to the AC for 48 hours. Finally, the neurons were fixed, and immunofluorescence for MAP2 was performed. **(I)** Representative images of the CB (red indicates Ctb-555) of compartmentalized neurons whose cell bodies were treated with DMSO (control) or LY294002 and whose axons were treated with or without BDNF. Scale bar = 20 µm. **(C-E)** Quantification of primary dendrites **(C)** and branching points **(D)** Sholl analysis **(E)** for neurons labeled with EGFP/MAP2/Ctb-555. n= 25-30 neurons from 3 independent experiments. *p<0.05, **p< 0.01, ***p< 0.001, +p<0.05 the LY294002 group vs. the LY294002/BDNF group. The results are expressed as the mean ± SEM. The data were analyzed by one-way ANOVA followed by Bonferroni’s multiple comparisons post hoc test (C, D, G, J, and K). Statistical analysis of the Sholl analysis data was performed by two-way ANOVA followed by Bonferroni’s multiple comparisons post hoc test.

Once PI3K is activated, it activates the mTOR kinase pathway by reducing the GAP activity of the Rheb GAP, leading to the activation of mTOR (Garza-Lombo and Gonsebatt, 2016). First, to evaluate BDNF-mediated activation of the mTOR pathway in cortical neurons, we stimulated noncompartmentalized cultures with BDNF for 1 hour in the presence or absence of LY294002 (a PI3K inhibitor) and Torin1, a specific inhibitor of mTOR that reduces the activity of both the TORC1 and TORC2 complexes (Liu et al., 2010). Western blotting revealed that BDNF promoted the phosphorylation of TrkB, Akt, S6r, and 4E-BP1 in cortical neurons and that LY294002 and Torin1 fully inhibited the phosphorylation of Akt, S6r and 4E-BP1 but not TrkB (Figure S4C), as expected. Then, we evaluated whether the mTOR pathway is involved in increasing protein translation upon arrival of pTrkB signaling endosomes in the cell body. To this end, we evaluated the time course of 4E-BP1 phosphorylation in the neuronal cell body induced by axonal BDNF stimulation. We observed that BDNF increased the phosphorylation of 4E-BP1 at 90 min and that 4E-BP1 phosphorylation was even higher at 180 min (Figure 7A and B). Similar results were obtained for the phosphorylation of S6r (Figure S4D and E). To evaluate whether activation of 4E-BP1 by axonal BDNF is dependent on the somatodendritic activity of PI3K and mTOR, we applied LY294002 or Torin1 to the CB of compartmentalized neurons (Figure 7C). The somatodendritic 4E-BP1 phosphorylation induced by axonal BDNF was significantly reduced by these inhibitors (Figure 7D and E). Additionally, treatment of axons with the dynein inhibitor CilioD reduced somatodendritic 4E-BP1 phosphorylation induced by axonal BDNF (Figure 7D and E). Together, these results indicate that upon arrival in the cell body, BDNF signaling endosomes induce activation of a signaling cascade that impacts mTOR-dependent protein synthesis.

**Figure 7.**
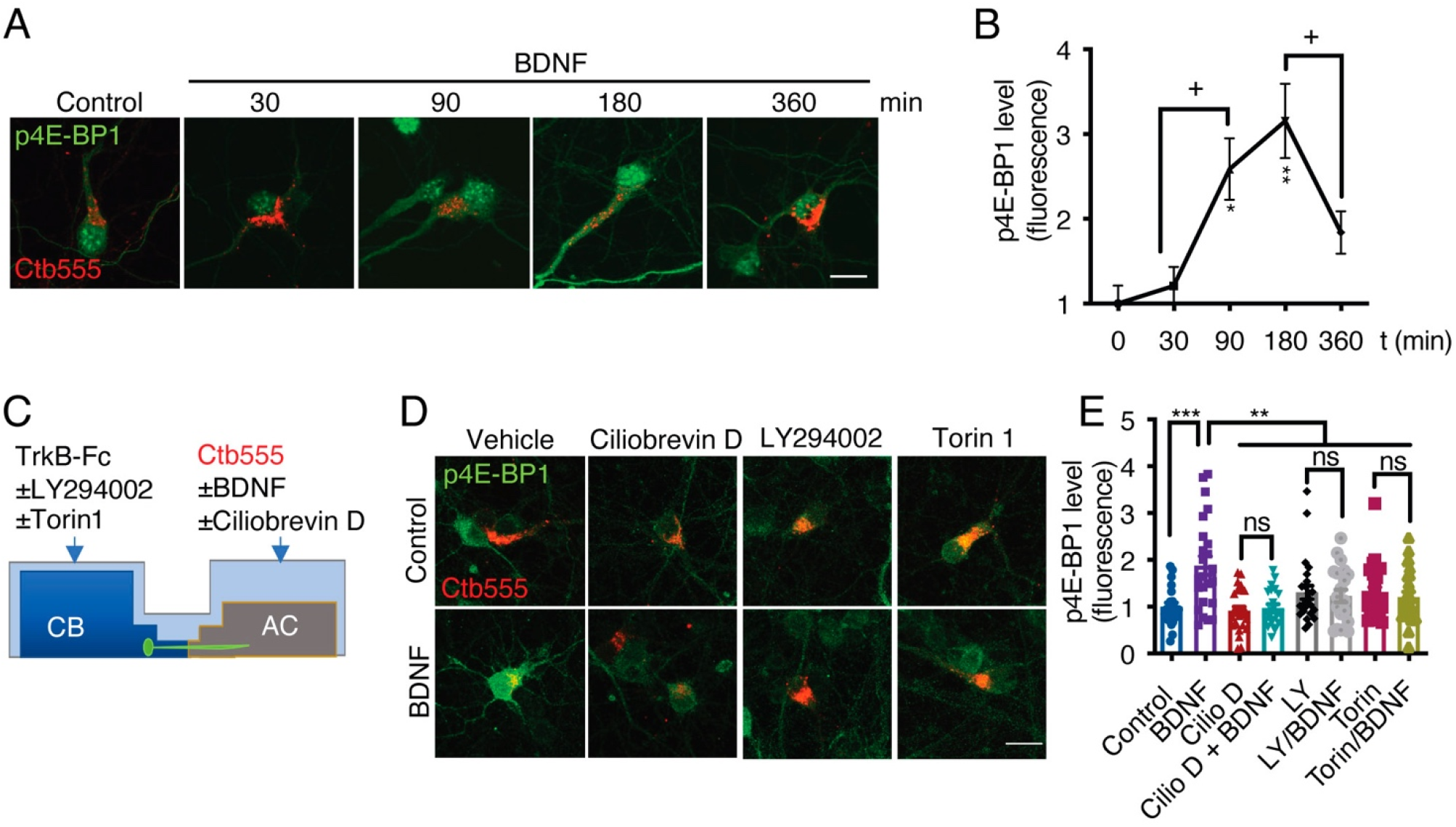
Axonal BDNF promotes mTOR activation in the cell bodies of compartmentalized cortical neurons in a PI3K-dependent manner. **(A)** Representative images of p4E-BP1 immunostaining (green) and Ctb-555 (red). DIV six compartmentalized cortical neurons were incubated with Ctb-555 overnight. At DIV 7, BDNF (50 ng/mL) was added to the AC of the neurons for 30, 60, 180 or 360 min. Scale bar = 20 µm**. (B)** Quantification of somatodendritic p4E-BP1 immunofluorescence in primary dendrites over time. **(C)** Schematic representation of the protocol used to evaluate the effect of different pharmacological inhibitors on the axonal BDNF-induced phosphorylation of 4E-BP1 in the cell body. DMSO (control), LY294002 (10 µm; LY), and Torin 1 (0.25 µm; Torin) were added to the CB of DIV six cortical neurons, or CilioD (20 µm; CilioD) was applied to the AC for 1 hour. Then, BDNF was added to the AC for 180 min in the presence or absence of these inhibitors. **(D)** Representative images of p4E-BP1 (green) in the somatodendritic compartment of neurons stimulated with BDNF in the presence or absence of different inhibitors. **(E)** Quantification of somatodendritic p4E-BP1 expression in neurons labeled with Ctb-555 (red) under each treatment. Scale bar = 20 µm. n= 31-36 neurons from 3 independent experiments. *p<0.05, **p< 0.01, ***p< 0.001. vs. the control group; +p<0.05 vs. the 90- and 360-min BDNF treatment groups in B. The results are expressed as the mean ± SEM. The data were analyzed by one-way ANOVA followed by Bonferroni’s multiple comparisons post hoc test.

### CREB and mTOR activity is required for protein synthesis in the somatodendritic compartment

The results presented above indicate that axonal BDNF induces the phosphorylation of CREB (Figure 3) and the activation of mTOR (Figure 7), suggesting that BDNF/TrkB signaling endosomes promote the translation of newly synthesized transcripts. To test this hypothesis, we metabolically labeled newly synthesized protein using Click-iT chemistry, which involves incorporation of L-azidohomoalanine (AHA). AHA is a modified amino acid that resembles methionine (Dieterich et al., 2010). In addition, we evaluated the level of *de novo* protein synthesis by using immunofluorescence to assess the levels of Arc, an immediate-early gene that encodes a protein required for synaptic plasticity induced by BDNF (Ying et al., 2002) and whose synthesis is regulated by CREB and mTOR (Ying et al., 2002, Takei et al., 2004). First, we adapted the Click-iT AHA method to noncompartmentalized neurons by stimulating the neurons with BDNF for 5 hours in the presence or absence of cycloheximide. As expected, BDNF increased AHA fluorescence, and cycloheximide significantly reduced the fluorescence of AHA (Figure S4F). We removed methionine and B27 from the medium of compartmentalized neurons for 1 hour. Next, we added AHA to the entire chamber and BDNF exclusively to the AC for 5 hours and treated the CB with or without KG501 or Torin1 CB (Figure 8A). As shown in Figure 8, BDNF significantly increased both AHA incorporation into newly synthesized proteins and Arc levels in the cell body and dendrites (Figure 8B-D). Remarkably, inhibition of CREB and mTOR resulted in significant reductions in BDNF-induced AHA incorporation into newly synthesized proteins and Arc protein expression in the cell body (Figure 8B-D). Together, these results indicate that long-distance BDNF/TrkB signaling induces the transcription and translation of specific mRNAs and that signaling endosomes play a role in coordinating both processes.

**Figure 8.**
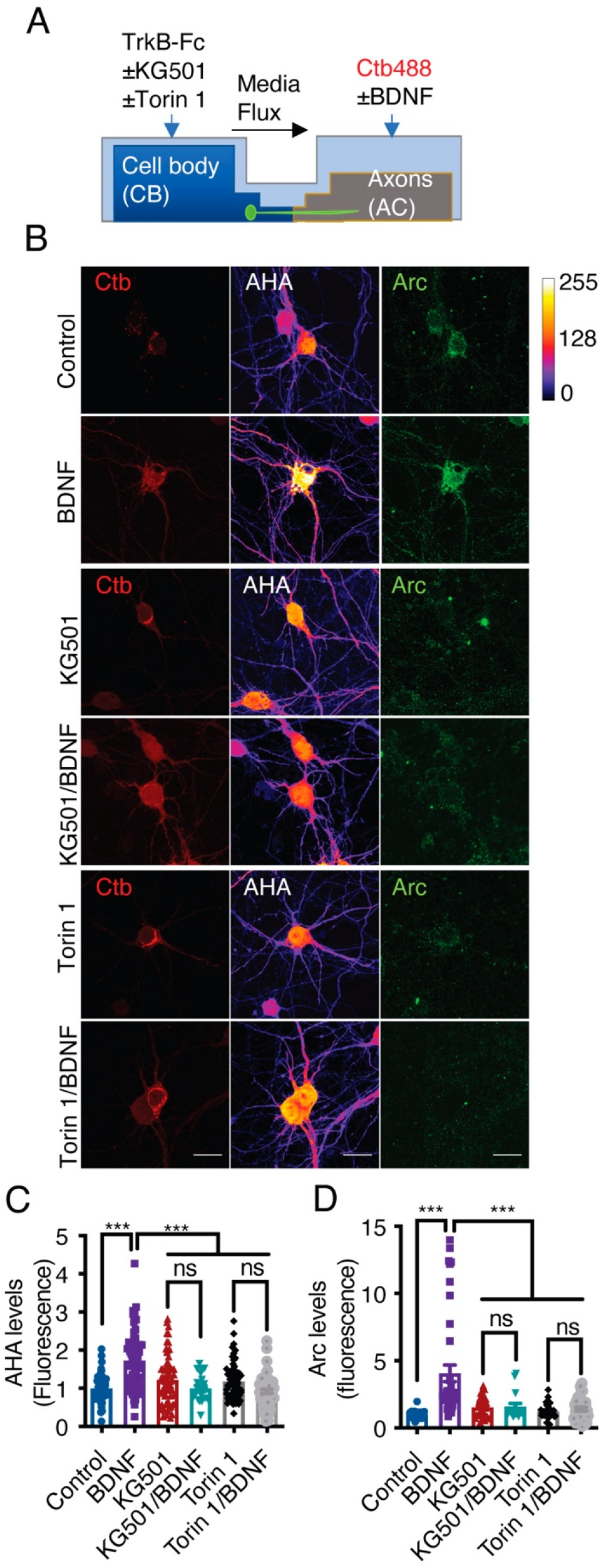
Axonal BDNF promotes somatodendritic protein synthesis in an mTOR- and CREB-dependent manner. **(A)** Schematic representation of the protocol used to evaluate protein synthesis. DIV 6-7 cortical neurons were incubated in methionine-free medium for 1 hour, and KG501 (10 µm) or Torin 1 (0.25 µm) was applied or not to the CB. Then, AHA was added to both the CB and AC, and BDNF (50 ng/mL) in the presence of Ctb-488 was added to only the AC for 5 hours. **(B)** Representative images of control and BDNF-stimulated neurons positive for Ctb-488 (red), AHA (in intensity color code mode, see color bar on the right) and Arc (green) in the presence or absence of KG501 and Torin 1. Scale bar = 20 µm **(C)** Quantification of AHA fluorescence in the primary dendrites of neurons labeled with Ctb-488 under each treatment. **(D)** Quantification of Arc fluorescence in the somas of neurons labeled with Ctb-488 under each different treatment. n= 30-48 neurons from 3 independent experiments. ***p< 0.001. The data were analyzed by one-way ANOVA followed by Bonferroni’s multiple comparisons post hoc test.

## DISCUSSION

In this study, we demonstrated, using microfluidic chambers and compartmentalized cultures derived from different mouse mutants, that BDNF/TrkB signaling endosomes are transported along the axons of cortical neurons in a dynein- and TrkB-dependent manner to regulate dendritic arborization. Although long-distance communication between neurotrophins in the PNS has been well described (Cosker and Segal, 2014, Scott-Solomon and Kuruvilla, 2018, Harrington and Ginty, 2013), whether signaling endosomes have a physiological role in the CNS in mammals is poorly understood. In this study, we clearly showed that BDNF/TrkB signaling endosomes originating in axons are required for dendritic arborization from the cell body induced by axonal BDNF. Our studies are in line with previous work showing that snapin, a protein that regulates autophagy and dynein transport, is required for dendritic branching in cortical neurons upon BDNF stimulation (Cheng et al., 2015)(Zheng et al., 2008, Zhou et al, 2012) and with the fact that BDNF/TrkB signaling endosomes generated in presynaptic boutons in the hippocampus locally regulate neurotransmitter release (Andres-Alonso et al., 2019). Additionally, our results are related to the retrograde effects of BDNF on dendritic arborization observed *in vivo* in retinal ganglion cells (Lom et al., 2002).

Using TrkB^F616A^ knock-in mice and 1NM-PP1 (Chen et al., 2005), we showed that the application of BDNF to axons increases the transport of activated TrkB to the cell body and that activated somatic TrkB in neurons is required for nuclear responses. The fact that 1NM-PP1 reduced the activation of already activated TrkB suggests that 1NM-PP1 binds the ATP binding site of the receptor, reducing the tyrosine activity of TrkB without affecting its neurotrophin binding (Chen et al, 2005). There are two possible mechanisms that can explain our results. The first is that TrkB is phosphorylated and dephosphorylated during its transport along the axon; thus, 1NM-PP1 binds the ATP binding cassette of the receptor and prevents its phosphorylation. The second possible mechanism is that BDNF retrograde signaling promotes the phosphorylation of somatodendritic TrkB and that this process is required to maintain CREB phosphorylation. In agreement with this last idea, other groups have shown that retrogradely transported TrkA in sympathetic neurons locally accumulates in the somatodendritic compartment, increasing the phosphorylation of other TrkA receptors in dendrites and the cell body (Lehigh et al., 2017, Yamashita et al., 2017).

Since BDNF can bind p75 and TrkB, we analyzed whether long-distance signaling is dependent on TrkB activation and the dependence of this process on p75 expression. p75 signaling has been reported to have an opposite role as Trk signaling since p75 can induce apoptosis or reduce axonal growth by binding to different ligands and receptors (Ibanez and Simi, 2012, Kraemer et al., 2014, Pathak et al., 2021). Additionally, p75 negatively regulates dendritic complexity and spine density in hippocampal neurons (Zagrebelsky et al., 2005). We observed that the absence of p75 resulted in a decrease in basal dendritic arborization of cortical neurons in microfluidic devices. Nevertheless, BDNF was able to rescue this phenotype, suggesting that p75 participates in the regulation of dendritic arborization but is not required for axonal BDNF signaling.

We found that PI3K activity in axons does not play a role in the retrograde transport of signaling endosomes. It was speculated that PI3K plays a role in this process due to the role of phosphoinositide accumulation in early endosome-mediated regulation of the early-to-late transition process (transition of Rab5- to Rab7-positive endosomes) (Numrich and Ungermann, 2014) required for retrograde sorting of signaling endosomes in motoneurons (Deinhardt et al., 2006). Our results are also different from findings regarding TrkA signaling in sympathetic neurons, as PI3K activity is required for the retrograde transport of TrkA signals (Kuruvilla et al., 2000). Nonetheless, the PI3K-mTOR signaling pathway is required for somatic responses to axonal BDNF. Indeed, our studies showed that PI3K activity is required for axonal BDNF-induced dendritic branching and phosphorylation of 4E-BP1 and S6r, which are downstream targets of mTOR, a process that is also dynein dependent. Consistent with these results, we demonstrated that axonal signaling endosomes activate the general translation of proteins in the cell body and that this process is dependent on CREB-mediated transcription and mTOR-dependent translation. These processes are accompanied by an increase in the protein levels of Arc. The fact that protein translation is also dependent on CREB suggests that CREB-dependent transcripts are specifically translated upon the arrival of signaling endosomes in the cell body in an mTOR-dependent fashion. *Arc* mRNA is anterogradely transported to dendrites to be locally synthesized (Steward and Worley, 2002), suggesting that axonal signaling may increase local translation in dendrites.

We performed a time-course study of CREB activation after application of BDNF to axons and found sustained activation of CREB after 3 hours of BDNF treatment. These results are different from what we have observed in noncompartmentalized hippocampal neurons, as CREB activity is decreased in these cells after one hour of BDNF treatment (Gonzalez-Gutierrez et al., 2020). While a dynein inhibitor reduced early activation of CREB by 40%, it reduced late CREB activation (after 3 hours of BDNF treatment) by 80%, suggesting that there is an additional component, most likely calcium waves, such as the ones reported in regenerating PNS neurons (Cho et al., 2013), that contributes to the early activation of CREB by axonal BDNF. Interestingly, it has been reported that both rapid and sustained CREB phosphorylation is required for proper transcriptional regulation (Dolmetsch et al., 2001), suggesting that in neurons, retrograde BDNF signaling may contribute to sustained CREB phosphorylation for CREB-dependent maintenance of dendritic arborization.

CREB plays a central role in processes such as learning and memory (Finkbeiner et al., 1997; Xing et al., 1998; Lonze and Ginty, 2002). Indeed, mutations in several coregulators or downstream TFs are associated with genetic diseases leading to autism and cognitive disabilities (Lyu et al, 2016; Zhou et al, 2016; McGirr et al, 2016; Wang et al, 2018). Several studies have suggested that axonal BDNF can activate CREB in the cell body (Zhou et al., 2012, Watson et al., 1999, Deinhardt et al., 2006, Bronfman et al., 2014); however, no studies have addressed the physiological role of CREB activation induced by axonal BDNF signaling. In this work, we showed that CREB is required for long-distance BDNF-induced dendritic arborization in cortical neurons, suggesting that axonal BDNF signaling may contribute to the development and maintenance of different circuits in the brain, including those involved in learning and memory in the hippocampus and motor learning in the cortico-callosal and cortico-striatal pathways (Andreska et al., 2020, Barco et al., 2005, Yap and Greenberg, 2018).

Several studies have shown that CREB is required for the development of dendrites *in vitro* or during development (Redmond et al., 2002, Kwon et al., 2011, Herzog et al., 2020, Gonzalez-Gutierrez et al., 2020). In our studies, we showed that as observed in mice with conditional deletion of TrkB in cortical neurons (Huang and Reichardt, 2003), reduced CREB activity was required for the dendritic maintenance of pyramidal cells. This is consistent with a report indicating that CREB is required for cortical circuit plasticity after stroke (Caracciolo et al., 2018).

Together, our results demonstrate that BDNF/TrkB signaling endosomes in axons can coordinate CREB-dependent transcription and mTOR-dependent protein translation in the cell body, likely wiring circuits in the CNS.

## MATERIALS AND METHODS

### Materials

See Supplementary Table 1 (Supplementary Material) for a list of all materials.

### Primary cortical neuron culture

Embryonic cortical neurons were obtained from mice (C57Bl/6J) and rats (*Rattus norvegicus*) (embryonic days 17–19) from the animal facilities of our institution. TrkB^F6161A^ knock-in mice (Ntrk2 ^tm1Ddg^/J), which have a single amino acid mutation in the intracellular ATP binding domain of TrkB that sensitizes the receptor to inhibition by 1NM-PP1 (Chen et al, 2005), and p75NTR−/− (B6.129S4-Ngfr^tm1Jae^/J) mice were purchased from The Jackson Laboratory (Sacramento, California, USA) and maintained as homozygotes (Ntrk2 ^tm1Ddg^/J) or heterozygotes (B6.129S4-Ngfr^tm1Jae^/J). Pregnant animals were euthanized under deep anesthesia according to bioethical protocols approved by the Bioethics Committee of the Pontificia Universidad Catolica de Chile and Universidad Andres Bello.

Rat and mouse cortical tissues were dissected out and dissociated into single cells in HBSS. After disaggregation, the neurons were resuspended in modified Eagle’s medium (MEM) supplemented with 10% horse serum (HS) (MEM/HS) and seeded in microfluidic chambers at a low density (40-50 x 10^3^ cells/chamber) or in mass culture at a density of 25 x 10^3^ cells/well on 12 mm coverslips or 1.8 x 10^6^ cells/60 mm plate. After four hours, the culture medium was replaced with neurobasal medium supplemented with 2% B27, 1x GlutaMAX and 1x penicillin/streptomycin. The proliferation of nonneuronal cells was limited by applying cytosine arabinoside (AraC; 0.25 µg/mL) when the MEM/HS was replaced with neurobasal medium and removed two days later (Shimada et al., 1998, Taylor et al., 2003).

### Microfluidic devices

The molds used to prepare the compartmentalized chambers were fabricated at the microfluidic core facility of Tel Aviv University (Gluska et al, 2016). The microfluidic chambers were prepared with a Syldgard^TM^ 184 silicone elastomer base according to the manufacturer’s instructions. Two days before primary culture, glass coverslips (25 mm) were incubated with poly-D-lysine (0.1 mg/mL). The next day, the poly-D-lysine was washed away, and microfluidic chambers with 400 µm microgrooves were placed on the coverslips. Then, laminin (2 µg/mL in water) was added to the chamber. On the same day of primary culture, the laminin solution was replaced with Dulbecco’s minimum essential medium (DMEM) supplemented with 10% HS, 1x GlutaMAX and 1x antibiotic/antimycotic (DMEM/HS).

### Quantification of BDNF-induced dendritic arborization in cortical neurons

For confocal microscopy analysis of cortical neurons in culture, a Nikon Eclipse C2 confocal microscope equipped with a digital camera connected to a computer with NIS-Elements C software was used. Images were acquired using a 60x objective at a resolution of 1024×1024 pixels along the z-axis unless otherwise indicated. Cortical neurons (DIV 6) were transfected with 0.5 µg of a plasmid expressing EGFP using 0.8 µL of Lipofectamine 2000 in 30 µL of Opti-MEM. After 2 hours, the Opti-MEM containing the plasmid was replaced with neurobasal media supplemented with 2% B27, 1x GlutaMAX, and 1x penicillin/streptomycin for 1 hour. Neurobasal medium supplemented with TrkB-Fc (100 ng/mL) was applied to the CB for all treatments. 1NM-PP1 (1 µM), KG501 (10 µM), and LY294002 (10 µM) were added to the CB, and K252a (0.2 µM), LY294002 (10 µM), and CilioD (20 µM) were applied to the AC. After 1 hour, BDNF (50 ng/mL) and fluorescently labeled Ctb (1 µg/mL) were added to the AC. After 48 hours, the neurons were washed with PBS at 37°C and then fixed with fresh 4% paraformaldehyde (PFA) in PBS (PFA-PBS) at 37°C for 15 min. Then, the chamber was removed, and the neurons were permeabilized, blocked with 5% BSA and 0.5% Triton X-100 in PBS, and incubated with anti-MAP2 (1:500) in incubation solution (3% BSA and 0.1% Triton X-100 in PBS). After being washed 3 times, the neurons were incubated with Alexa 647-conjugated donkey anti-mouse (1:500) in incubation solution and mounted for analysis by fluorescence microscopy using Mowiol 4-88.

Dendritic arborization in cortical neurons labeled with Ctb, transfected with EGFP and labeled with MAP2 was analyzed. Primary dendrites and branching points were quantified, and Sholl analysis (Sholl, 1953) was performed (see below).

### Stereotaxic brain surgery in mice

Two-month-old male C57BL76J mice were deeply anesthetized using a mixture of ketamine/xylazine (150 mg/kg, 15 mg/kg) and placed in a stereotaxic frame (RWD, Life Science Co.). AAV1 expressing CREB-DN (Gonzalez-Gutierrez et al., 2020) in frame with EGFP or EGFP alone (control) driven by the synapsin promoter and AAV1 expressing mCherry were coinjected unilaterally into II/III layer of the motor cortex at the following coordinates: +0.6 mm anteroposterior to bregma, -1.5 mm lateral bregma and -0.6 mm deep (Watakabe et al., 2014). The brains of control mice were coinjected with AAV1-hSyn-EGFP and AAV1-hSyn-mCherry (0.5 µL of each and 1 µL of 0.9% NaCl), and the brains of experimental mice were coinjected with AAV1-hSyn-EGFP-CREN-DN-WPRE-SV40Pa and AAV1-hSyn-mCherry (0.5 µL of each and 1 µL of 0.9% NaCl). 1×10^8^ PFU of each AAV1 was administered. To analyze dendritic arborization, the animals were sacrificed with a mixture of ketamine/xylazine (200 mg/kg, 20 mg/kg) in saline 3 weeks postinjection and perfused transcardially with 0.9% NaCl, followed by fixation with 4% PFA in phosphate buffer (pH 7.4). The mouse brains were dehydrated in 30% sucrose and sectioned at a thickness of 40 μm with a cryostat (Leica Microsystem), and the brain sections were immunostained for mCherry (1:1000).

### Analysis of dendritic arborization

Confocal images of cultured cells were acquired using a 60x objective at a resolution of 1024×1024 pixels along the z-axis. A total of 5-7 optical sections over a width of 0.5 covering the whole neuron were obtained. Images of tissue sections were acquired with a confocal laser microscope (Leica TCS SP8) with a 40x oil objective at a resolution of 1024×1024 pixels. Fifteen to 20 optical sections of isolated mCherry-labeled neurons in layer II/III of the sensorimotor cortex of injected mice were obtained over a width of 0.5 µm along the z-axis. Subsequently, the z-stacks were integrated, and neurons were manually drawn guided by the original fluorescence image using the pencil tool in ImageJ and segmented to obtain binary images. The numbers of total primary dendrites and branching points of all dendrites were manually counted from the segmented images using the Fiji open source platform (Schindelin et al., 2012). For Sholl analysis, concentric circles with increasing diameters (10 µm per step for primary cultured cortical neurons and 5 µm for cortical neurons in tissue sections) were traced around the cell body, and the number of intersections between dendrites and circles was counted and plotted for each circle. The analysis was performed using Sholl analysis plug in in Fiji (Ferreira et al., 2014). In addition, for cortical neurons in tissue sections, the cell body size and apical dendrite diameter were measured from the segmented images using the straight tool in ImageJ. Additionally, the width and length of the cell body and apical diameter were measured 5, 25 and 50 µm from the start of the apical dendrite.

### Evaluation of protein phosphorylation by immunofluorescence

Cortical neurons (DIV 5-6) were incubated with Ctb-555 (Ctb, 1 µg/mL) overnight. For all treatments, neurobasal medium supplemented with TrkB-Fc (100 ng/mL) was added to the CB of DIV 6-7 cortical neurons. LY294002 (10 µM) and Torin 1 (0.25 µM) were added to the CB, and CilioD (20 µM) was applied to the AC. After 1 hour, BDNF (50 ng/mL) was added to the AC. After the time point indicated in the figure, the samples were fixed with 4% PFA and phosphatase inhibitor in PBS for 15 min. The samples were blocked and permeabilized in blocking solution (5% BSA, 0.3% Triton X-100, and 1x phosphatase inhibitor in PBS) for 1 hour and incubated with antibodies overnight (4°C) in incubation buffer (3% BSA and 0.1% Triton X-100 in PBS). The following antibodies were used: anti-pCREB (1:500), anti-pS6r (1:100), and anti-p4E-BP1 (1:500). The samples were incubated with Alexa 488-conjugated donkey anti-rabbit secondary antibody (1:500) for 90 min in BSA (3%) and Triton X-100 (0.1%) in PBS and then incubated with Hoechst 33342 (5 µg/mL) to visualize the nuclei. Entire neurons in the CB were visualized by confocal microscopy, and five to seven optical sections with a thickness of 0.5 µm thick from a single cell were analyzed.

### Evaluation of neuronal viability by TUNEL staining

For the detection of fragmented DNA, KG501 (10 µM) was applied to the CB and Ctb-555 (1 µg/mL) was added to the AC of DIV 6 cortical neurons cultured in neurobasal medium supplemented with B27 for 48 hours. Oligomycin A (10 µM) was added to the CB for 5 min as a positive control. The neurons were washed with PBS and fixed in 4% PFA in PBS for 15 min at room temperature. After that, coverslips to which neurons were attached were washed with PBS three times for 5 min each. The neurons were permeabilized in□ice_cold citrate/Triton solution (0.1% citrate and 0.1% Triton X□100) for 2 min on ice. Then, the coverslips were washed twice to remove the permeabilization solution and incubated with TUNEL labeling mixture (In Situ Cell Death Detection Kit) for 1 hour at 37°C. The samples were incubated with Hoechst (5 µg/mL). TUNEL staining was analyzed by confocal microscopy, and Hoechst-positive nuclei and TUNEL-positive apoptotic nuclei were counted by using the Fiji platform.

### Evaluation of retrograde transport of TrkB from axons to cell bodies via immunoendocytosis of Flag-TrkB in microfluidic chambers

Cortical neurons (DIV 5) were transfected with 0.5 µg of plasmid expressing Flag-TrkB (gift from Prof. Francis Lee, NYU, USA) using 0.8 µL of Lipofectamine 2000 in 30 µL Opti-MEM per chamber. After 48 hours, cortical neurons (DIV 7) were incubated at 4°C for 10 min, and then an anti-Flag antibody (1:750) was applied to the AC for 45 min at 4°C. Then, neurons were briefly washed with warm neurobasal medium, and BDNF (50 ng/mL) was added to the AC for 180 min. Finally, the samples were fixed with 4% PFA and a phosphatase inhibitor in PBS for 15 min. The samples were blocked and permeabilized in blocking solution (5% BSA, 0.3% Triton X-100, and 1x phosphatase inhibitor in PBS) for 1 hour and incubated with an anti-pTrkB (Y816) (1:200) or anti-Rab5 (1:250) antibody in incubation buffer (3% BSA and 0.1% Triton X-100 in PBS) overnight (4°C). The samples were incubated with Alexa 488-conjugated donkey anti-rabbit (1:500) and Alexa 555-conjugated donkey anti-mouse (1:500) secondary antibodies for 90 min in incubation buffer. We visualized entire neurons in the CB and the microgroove compartment via confocal microscopy using a confocal microscope. Five to seven optical slices with a thickness of 0.5 µm were analyzed for each cell.

### Evaluation of somatodendritic TrkB activity in cortical neurons from TrkB^F616A^ mice in compartmentalized cultures

Ctb-555 (1 µg/mL) was applied to the AC of cortical neurons from TrkB^F616A^ mice (DIV 6) overnight. Then, the neurons were washed with neurobasal medium. For control and BDNF treatment, neurobasal medium (without B27) supplemented with TrkB-Fc (100 ng/mL) was added to the CB, and the AC was treated with neurobasal medium with and without BDNF (50 ng/mL) for 3 hours. To inhibit TrkB activity in the CB, the culture medium was removed from the CB and 1NM-PP1 (1 µM) was added to the CB for one hour; additional medium was added to the AC to avoid leakage of 1NM-PP1 into the AC. Next, the AC of the neurons was treated with or without BDNF (50 ng/mL), with additional medium being added to the AC, and 1NM-PP1 (1 µM) and TrkB-Fc (100 ng/mL) were applied to the CB for 3 hours. When the activity of TrkB in the AC was inhibited, 1NM-PP1 (1 µM) was added to the AC for 1 hour. Then, BDNF (50 ng/mL) was applied to the AC in the presence of 1NM-PP1 (in AC) and TrkB-Fc (in CB) for 3 hours, with additional medium being added to the CB. In all conditions, TrkB-Fc (100 ng/mL) in neurobasal medium without B27 was added to the CB. After 3 hours of BDNF treatment, the neurons were washed with 1x PBS, and the cells were fixed with 4% PFA in PBS for 15 min at room temperature. Finally, immunofluorescence for phosphoproteins was performed as described above. The expression of pTrkB (Y816; 1:200) and pCREB (S133, 1:500) was evaluated. To assess pTrkB and pCREB levels, we visualized the CB of entire neurons by confocal microscopy. The number of pTrkB puncta in the cell body were quantified in Ctb-positive cells and normalized to the size of the cell body. pCREB levels were evaluated as described above.

### Evaluation of TrkB activity and downstream signaling in noncompartmentalized cortical neurons from TrkB^F616A^ mice

Cortical neurons from TrkB^F616A^ mice (DIV 7) were treated with or without 1NM-PP1 (1 µM) in neurobasal medium (without B27) for 1 hour. Then, control neurons were treated with vehicle for 1 hour; neurons in the treatment group were incubated with BDNF (50 ng/mL) for 30 min, rinsed twice with 1x PBS and incubated with vehicle for 30 min; neurons in the 1NM-PP1 (Pre) group were incubated with BDNF in the presence of 1NM-PP1 for 1 hour; and neurons in the 1NM-PP1 (Post) group were stimulated with BDNF for 30 min. Next, the cells were rinsed twice with 1x PBS and incubated with 1NM-PP1 for 30 min. Finally, TrkB and Akt activity was evaluated by Western blotting, and CREB, 4E-BP1 and S6r expression was evaluated by immunofluorescence. For western blotting, the cells were lysed with RIPA buffer (0.1% SDS, 0.5% NP40, 10 mM Tris-HCl (pH 7.5), 1 mM EDTA, 150 mM NaCl, and 0.5% deoxycholic acid) containing protease and phosphatase inhibitors. The cell extracts were subjected to standard SDS gel electrophoresis and Western blotting protocols using anti-pTrkB (Y816, 1:1000), TrkB (1:1000), pAkt (1:1000), Akt (1:1000), and GAPDH (1:1000) antibodies. For immunofluorescence, the neurons were washed with 1x PBS and fixed with 4% PFA in PBS for 15 min at room temperature. Finally, immunofluorescence for phosphorylated proteins was performed as described above using the following antibodies: anti-pCREB (S133; 1:500), anti-pS6r (S235/236, 1:100), anti-p4E-BP1 (S65, 1:500) and anti-β-III tubulin (1:750). We visualized the CB of neurons by confocal microscopy. Images were acquired using a 20x objective for pCREB and a 60x objective for 4E-BP1 and S6r at a resolution of 1024×1024 pixels along the z-axis of whole cells.

### Evaluation of protein synthesis by Click-iT chemistry and Arc immunofluorescence

Ctb-488 was added to the AC of compartmentalized cortical neurons (6 DIV) overnight. At DIV 7, the neurons were washed with warm 1x PBS and incubated with DMEM without methionine supplemented with GlutaMAX 1x and L-cystine for 60 min in the presence or absence of Torin 1 (0.25 µM) or KG501 (10 µM). The medium was replaced with DMEM supplemented with Click-iT AHA (0.1 mM) for 5 hours. The AC of neurons was treated with BDNF (50 ng/mL), and the CB compartment was incubated with TrkB-Fc (100 ng/mL) and Torin 1 (0.25 µM) or KG501 (10 µM). The neurons were fixed with 4% PFA in PBS for 15 min. Then, alkyne-Alexa Fluor 555 (2.5 µM) was conjugated to AHA according to the manufacturer’s instruction. Finally, immunofluorescence for Arc (1:300) was performed as described above.

### BDNF-avi production, purification, biotinylation and Q-Dot conjugation

Production of BDNF-QDs was performed according to (Stuardo et al., 2020). Briefly, HEK293FT cells were cultured in DMEM, 4.5 g/l glucose, 10% FBS, and 1% penicillin/streptomycin. For transfection, HEK293FT cells were grown in 20 plates of 15 cm until they reached ∼70% confluency. The medium was replaced with 25 ml of high-glucose serum-free DMEM. For transfection, 30 µg of pcDNA3.1-BDNFavi (gift from Prof. Chengbiao Wu, UCSD, USA) was added to one ml of high-glucose DMEM, and then 90 µg of PEI (45 µL) was added. The mixture was incubated at room temperature for 25 min and added dropwise to the medium. Transfected HEK293FT cells were incubated at 37°C in 5% CO_2_ for 48 hours, and then the medium was collected for protein purification.

The medium was harvested and treated with 30 mM phosphate buffer (pH 8.0), 500 mM NaCl, 20 mM imidazole, and protease inhibitor cocktail. After incubation on ice for 15 min, the medium was cleared by centrifugation at 9500 rpm for 45 min using a Hettich 46R centrifuge. Ni-NTA resin was rinsed twice with washing buffer (30 mM phosphate buffer (pH 8.0), 500 mM NaCl, 20 mM imidazole, and protease inhibitor cocktail) and added to the medium at a concentration of 0.3 ml Ni-NTA/100 ml media. Then, the samples were incubated for 180 min at 4°C. The medium/Ni-NTA slurry was loaded onto a column, and the captured Ni-NTA resin was washed with 10 ml wash buffer and eluted with 1 ml elution buffer (30 mM phosphate buffer (pH 8.0), 500 mM NaCl, 300 mM imidazole, and protease inhibitors). The purity and concentration of BDNF-avi were assessed by Western blotting using a standard BDNF curve and an anti-BDNF antibody as done previously (Stuardo et al., 2020).

BDNF-avi was dialyzed against Sanderson water for 15 min. Then, BDNF-avi protein was biotinylated via an enzymatic reaction using the enzyme BirA. BDNF-avi (600-800 ng) was incubated with biotinylation buffer (50 mM bicine, 10 mM MgOAc, and 50 µM D-biotin) in the presence of 10 mM ATP and BirA-GST (BirA 1:1 BDNF-avi) for 1 hour at 30°C with agitation at 600 rpm. Then, 10 mM ATP and BirA-GST were added again, and the samples were incubated for an additional hour at 30°C with agitation at 600 rpm.

For coupling of BDNF-bt to streptavidin QDs and visualization in microfluidic devices, 1 µ L of QD-655 was added to 10 µL of BDNF-bt (approx. 3 ng/µL) to allow the binding of BDNF-bt to streptavidin QD-655. Then, neurobasal medium was added to the BDNF-QDs to a final volume of 20 µL, and the mixture was incubated at room temperature in an orbital rocker for 30 min. To evaluate the role of PI3K and dynein in BDNF-QD transport along axons, maintenance medium was replaced with neurobasal medium without B27, and AC of neurons were treated for 1 hour in the presence or absence of LY294002 (10 µM) or CilioD (20 µM), respectively. Then, BDNF-QDs (2 nM) were added to the AC in the presence or absence of LY294002 or CilioD for 4 hours. The neurons were fixed with 3% PFA in PBS for 15 min at room temperature, washed with PBS, and mounted in Mowiol. We visualized the proximal part of the microgrooves in the CB by confocal microscopy. Three to five optical sections (3-5) with a thickness of 0,25 µm were analyzed for each axon. Images were acquired using a 100x objective at a resolution of 1024×1024 pixels along the z-axis. The number of BDNF-QDs or QDs alone was quantified in 90 µm^2^ regions of axons located in the microgrooves.

### Statistical analysis

The results are expressed as the average ± standard error of the mean (SEM). Sholl analysis curves were analyzed by two-way repeated-measures ANOVA followed by Bonferroni’s multiple comparisons test. Student’s t test or one-way ANOVA followed by an appropriate multiple comparisons test was performed depending on the number of groups in the experiment. Details about the specific test used, level of significance, and number of replicates are indicated in each figure legend. Statistical analyses were performed using GraphPad Prism 7 (Scientific Software).

## Acknowledgments

The authors thank Carolina Ramirez for help with the compartmentalized cultures, Nicolas Stuardo for help with the BDNF monobiotinylation protocol and Carlos Ibañez for helpful reading of the manuscript.

---

**Figure S1.**
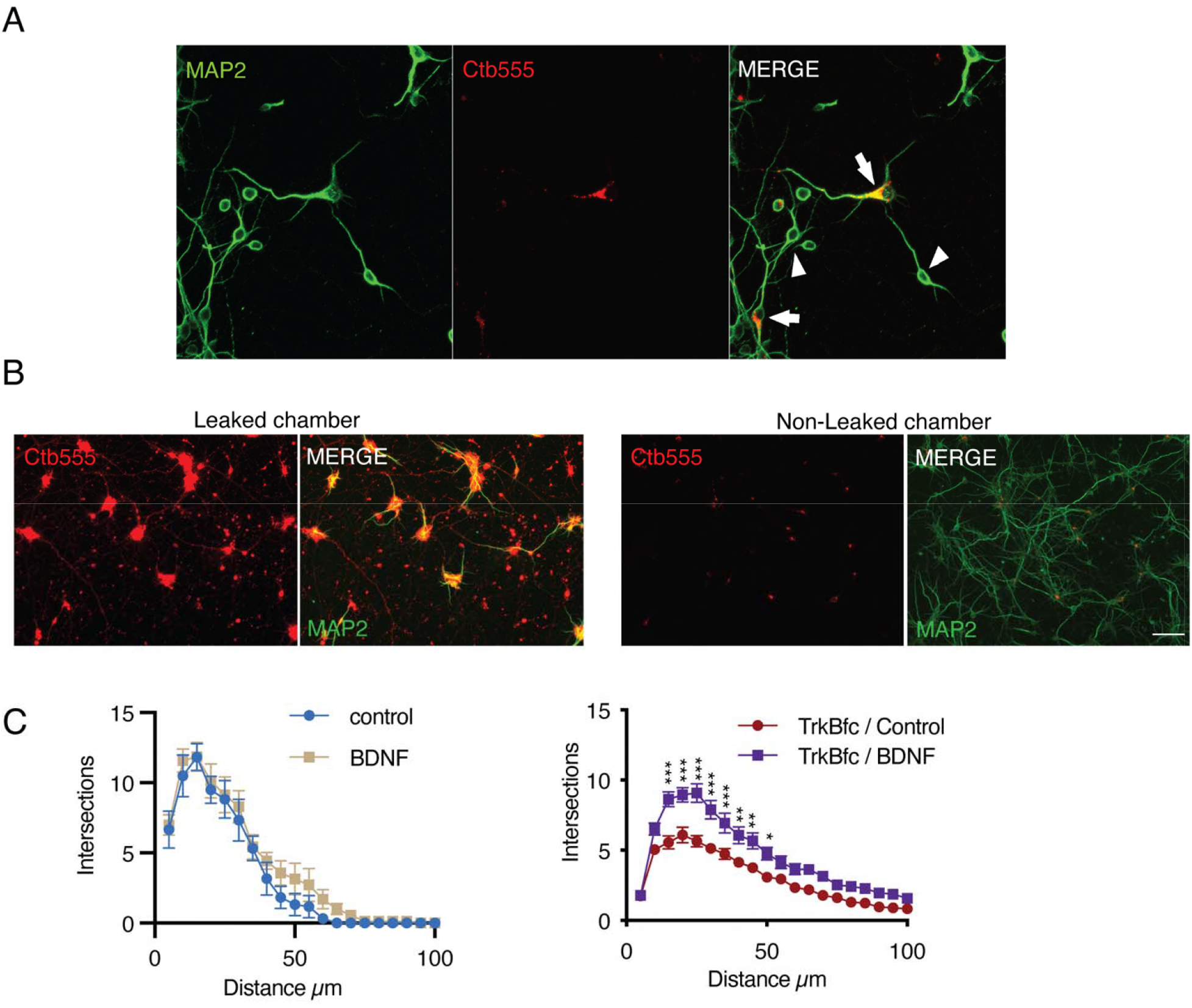
Experimental design used to study retrograde BDNF signaling in compartmentalized cortical neurons. **(A)** Representative images of compartmentalized neurons whose axons were incubated with Ctb-555 for 48 hours. Neurons were immunostained for MAP2 (green). The arrows indicate neurons that projected axons into the AC. The arrowheads indicate neurons with axons that do not project to the AC and therefore are not labeled with Ctb-555. **(B)** Representative images of a microfluidic chamber that were not well compartmentalized (leaking, left panel) and one in which compartmentalization was achieved (no leaking, right panel). Ctb-555 is visualized in red, and MAP2 is visualized in green. Scale bar = 50 μm. (C) Sholl analysis of dendritic arborization of compartmentalized rat cortical neurons stained with Ctb-555. Left panel, BDNF (50 ng/mL) was applied or not to the AC for 48 hours. Right panel, TrkB-Fc (100 ng/mL) was added to the CB (n=10-15 neurons, from 2 independent experiment), and BDNF (50 ng/mL) was applied or not to the AC for 48 hours (n=37 neurons, from 2 independent experiment). The results are expressed as the mean ± SEM. ***p<0.001. The Sholl analysis data were analyzed by two-way ANOVA followed by Bonferroni’s multiple comparisons post hoc test.

**Figure S2.**
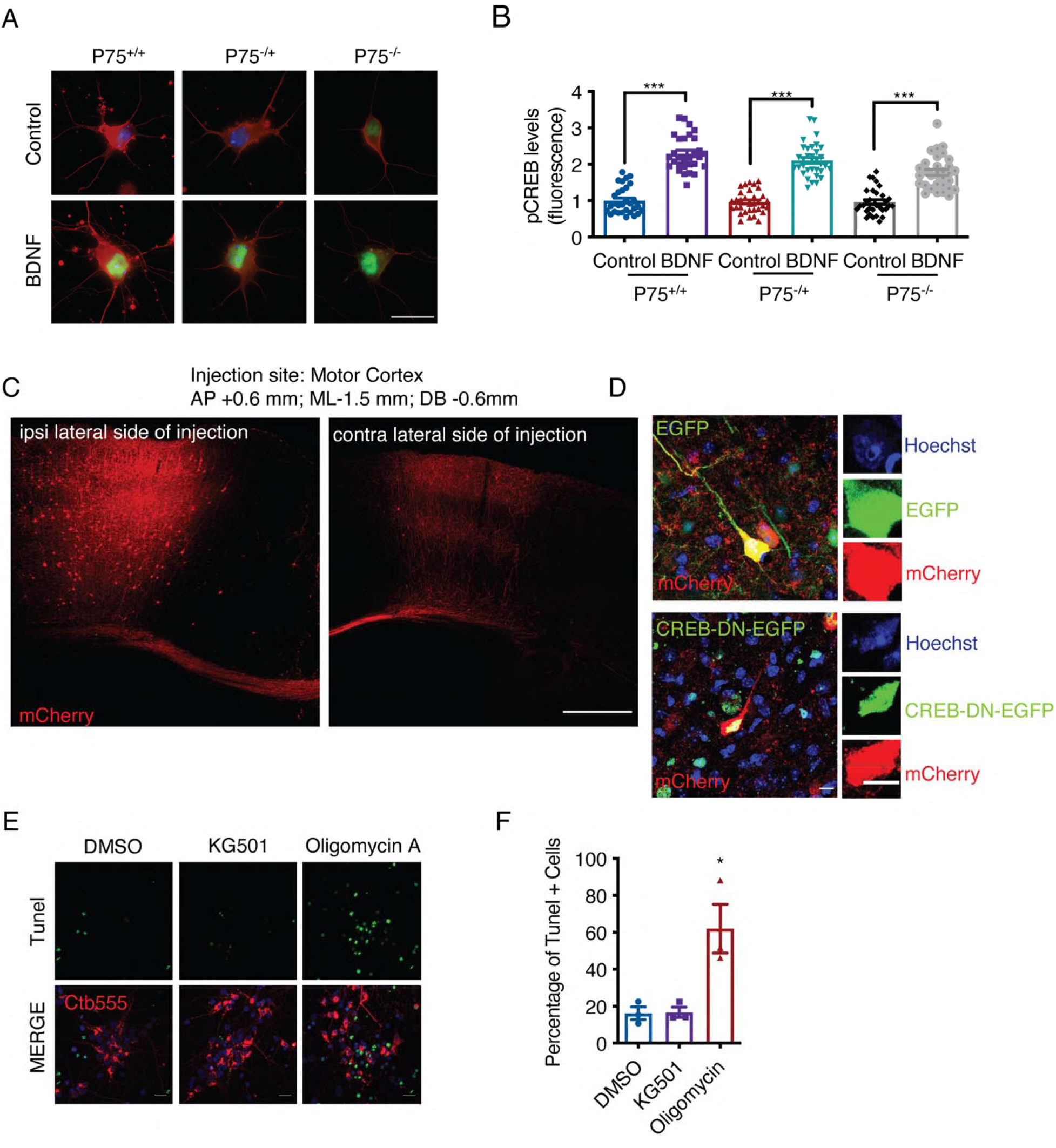
Evaluation of neuronal survival and CREB activation. DIV seven cortical neurons from p75^+/+^, p75^+/-^ or p75^-/-^ mice were treated with BDNF (50 ng/mL) for 30 min, washed and fixed for immunostaining for pCREB (S133), as indicated in the methods section. **(A)** Representative image of pCREB (green) and βIII-tubulin (red) in BDNF-stimulated neurons from mice of each p75 genotype. Scale bar = 20 µm. **(B)** Quantification of the pCREB fluorescence intensity in the nuclei of neurons derived from mice of the 3 different p75 genotypes treated with or without BDNF. n= 30 neurons from 2 independent experiments. The results are expressed as the mean ± SEM. ***p< 0.01. The data were analyzed by one-way ANOVA followed by Bonferroni’s multiple comparisons post hoc test. **(C)** Representative image of a mouse brain injected unilaterally with AAV into layer II/III of the motor cortex. Scale bar = 500 µm **(D)** Representative image of neurons expressing EGFP/mCherry or CREB-DN-EGFP/mCherry. Right panel, Hoechst staining was used to visualize nuclear morphology. **(E)** To control cell death induced by KG501, we performed the experiments described in D after visualizing DNA fragmentation in fixed cells by TUNEL staining (green). Only neurons labeled with Ctb-555 were analyzed. The neurons were treated with oligomycin (10 μM) as a positive control as indicated in the methods section. **(J)** Quantification of TUNEL-positive cells. Scale bar =20 μm. n=2 chambers per condition from 3 independent experiments. *p=0.05. The results are expressed as the mean ± SEM. The data were analyzed by one-way ANOVA followed by Bonferroni’s multiple comparisons post hoc test.

**Figure S3.**
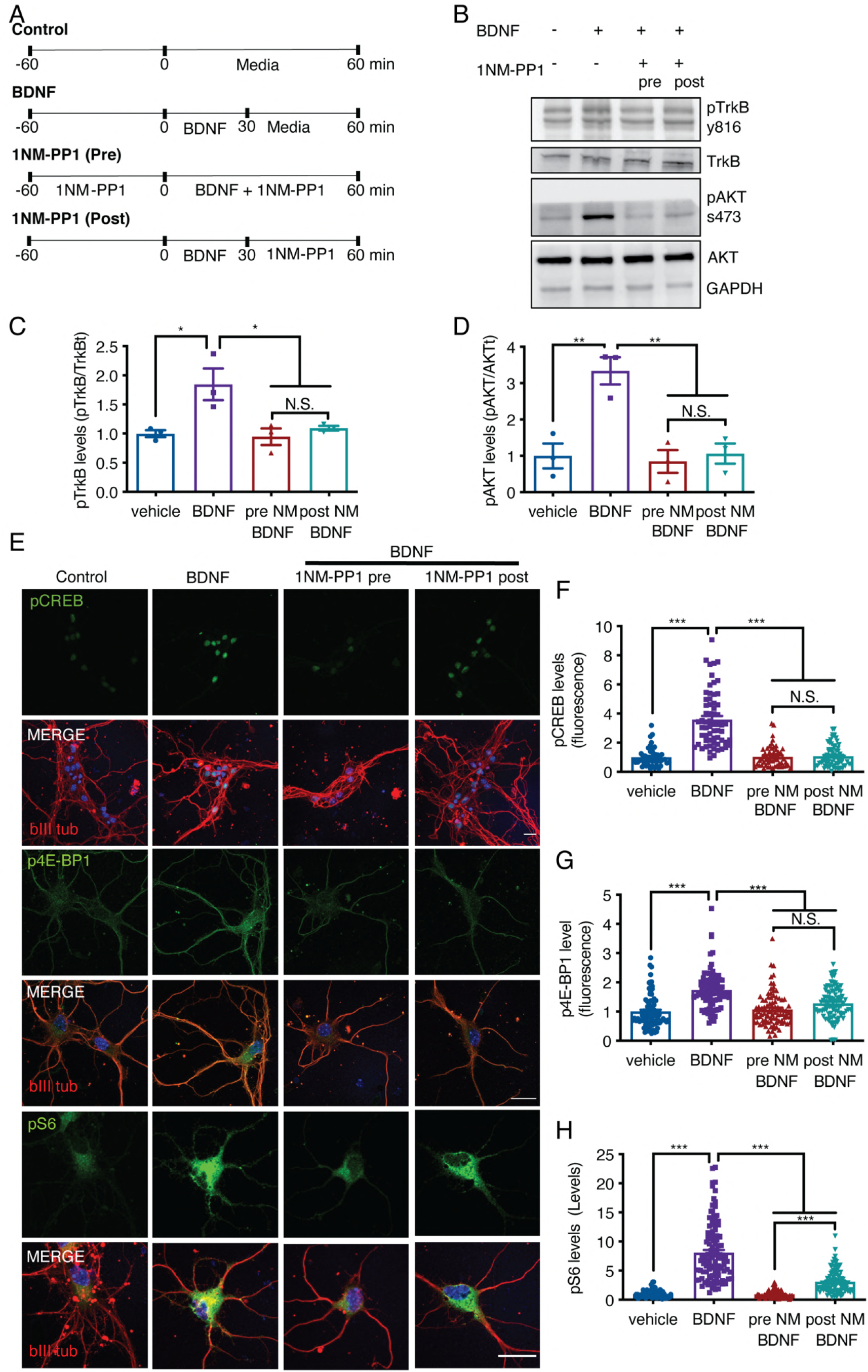
1NM-PP1 can reduce TrkB activation in TrkBF616A mouse cortical neurons after BDNF treatment. **(A)** Diagrams of the experimental designs used to stimulate neurons in noncompartmentalized cultures. The medium of DIV 7 cortical neurons was removed for 1 hour, and the neurons were treated with or without 1NM-PP1 (1 μM). Then, the neurons were treated as follows: neurons in the control group were treated with vehicle for 1 hour; neurons in the BDNF group were incubated with BDNF (50 ng/mL) for 30 min, rinsed twice and incubated with vehicle; neurons in the 1NM-PP1 (Pre) group were incubated with BDNF in the presence of 1NM-PP1 for 1 hour; and neurons in the 1NM-PP1 (Post) group were incubated with BDNF for 30 min, rinsed twice and incubated with 1NM-PP1 for 30 min. **(B)** Immunoblotting of pTrkB (Y816), TrkB, pAkt (S437), Akt and GAPDH in cortical neurons stimulated as described in A. (**C**) Quantification of pTrkB levels normalized to total TrkB levels. **(D)** Quantification of pAkt levels normalized to total Akt levels. n=3 independent experiments. **(E)** Representative image of cortical neurons treated as described in A. pCREB, p4E-BP1 and pS6 are visualized in green, and βIII tubulin is shown in red. Scale bar = 20 μm**. (F)** Quantification of pCREB fluorescence (arbitrary units, A.U.) in the nuclei of neurons. **(G)** Quantification of p4E-BP1 fluorescence (arbitrary units, A.U.) in primary dendrites. **(H)** Quantification of pS6 fluorescence (arbitrary units, A.U.) in primary dendrites. n=49-72 neurons from 3 independent experiments. The results are expressed as the mean ± SEM. *p< 0.05, **p<0.01, ***p< 0.001. The data were analyzed by one-way ANOVA followed by Bonferroni’s multiple comparisons post hoc test.

**Figure S4.**
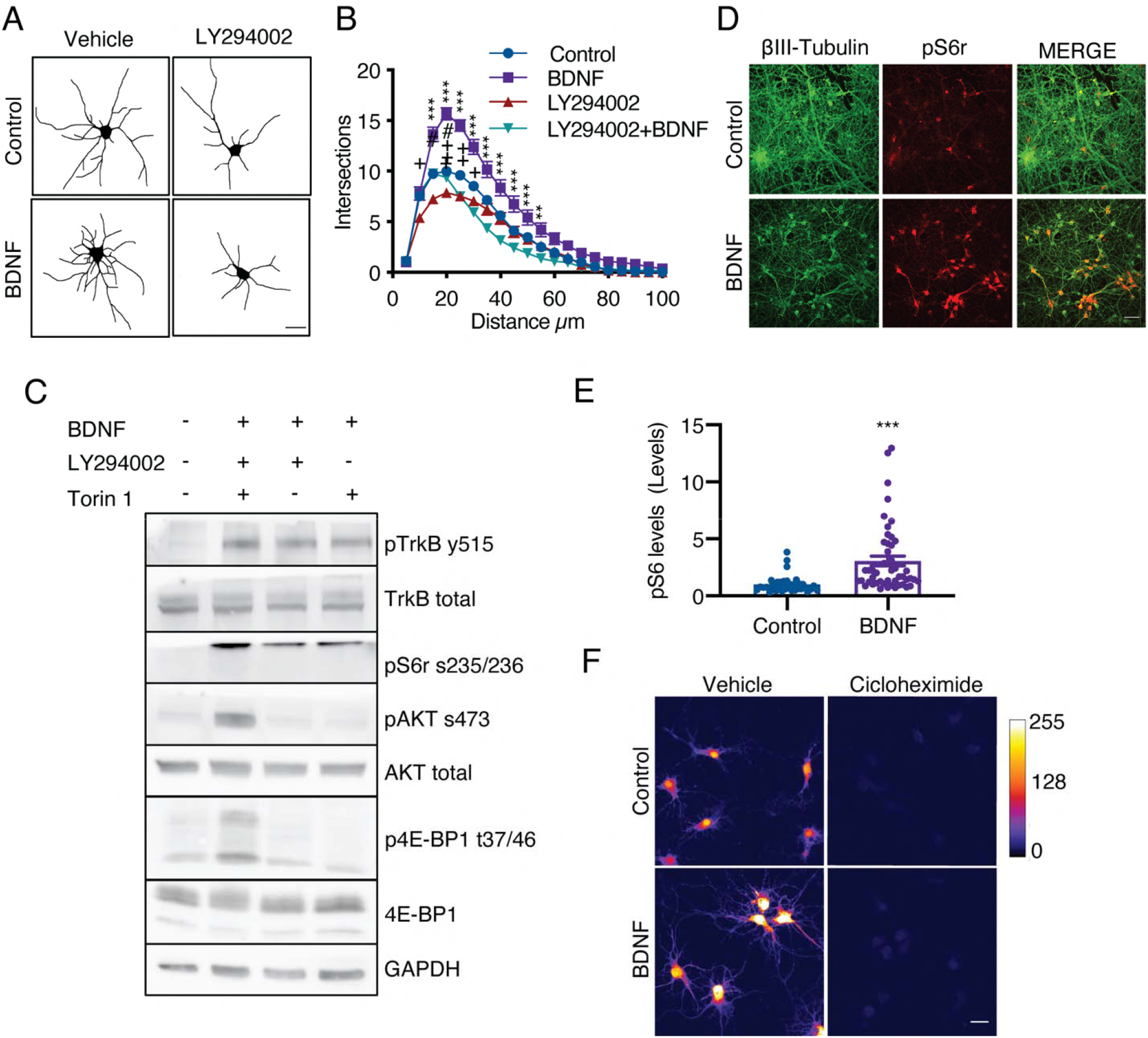
PI3K signaling is required for BDNF signaling in cortical neurons. **(A)** Representative images of rat cortical neurons in non-compartmentalized cultures treated with vehicle, LY294002 (10 M), BDNF (50 ng/mL) or BDNF after preincubation with LY294002. Scale bar = 20 µm. **(B)** Quantification of total dendritic branching by Sholl analysis. n=25-32 from 2 independent experiments. **p<0.01 and ***p<0.001, the control group vs. the BDNF group; ++p<0.01, the control group vs. the LY294002 group. The results are expressed as th mean ± SEM. The Sholl analysis data were analyzed by two-way ANOVA followed by Bonferroni’s multiple comparisons post hoc test. **(C)** Western blot analysis of DIV 7 non-compartentalized neurons treated with BDNF (50 ng/mL) in the presence or absence of LY294002 (10 μM) or Torin 1 (0.25 μM) for 1 hour. pTrkB (Y515), TrkB, pAkt (S473), total Akt, pS6r (S235/236), p4E-BP1 (T37/46), total 4E-BP1 and GAPDH expression was evaluated. **(D)** Representative images of β-III-tubulin (green) and pS6r (red) in cortical neurons whose axons were stimulated with BDNF (50 ng/mL) for 180 min. Scale bar = 50 μm. (**E**) Quantification of pS6r fluorescence intensity in the soma of neurons treated with or without BDNF. n= 36-48 neurons from 3 independent experiments. The results are expressed as the mean ± SEM. ***p< 0.01. Student’s t-test was used for statistical analysis. **(F)** Representative image of AHA staining in neurons treated with BDNF for 5 hours in the presence or absence of cycloheximide. Right panel, color code for intensity. Scale bar = 20 µm.

**Supplementary Table 1.**
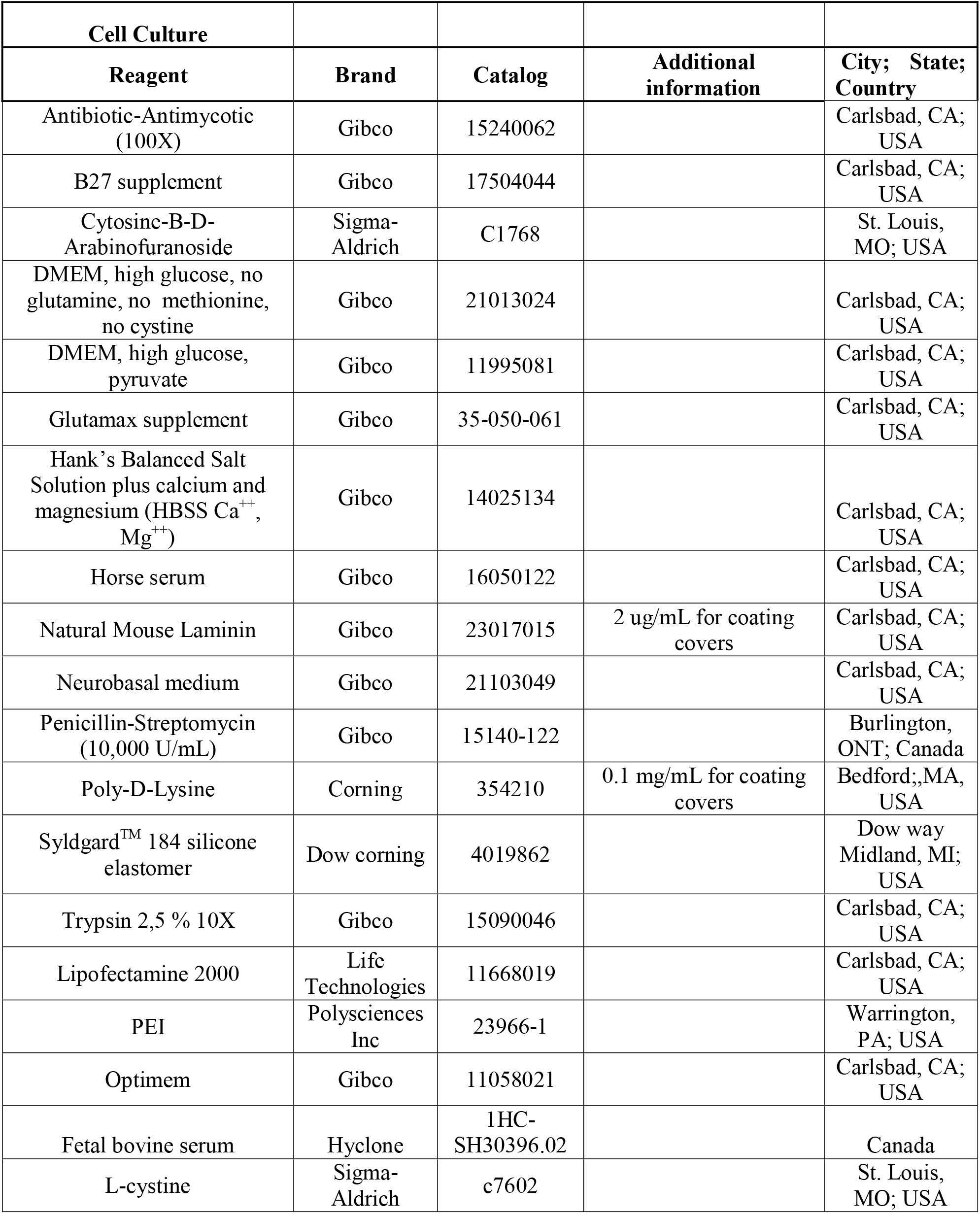
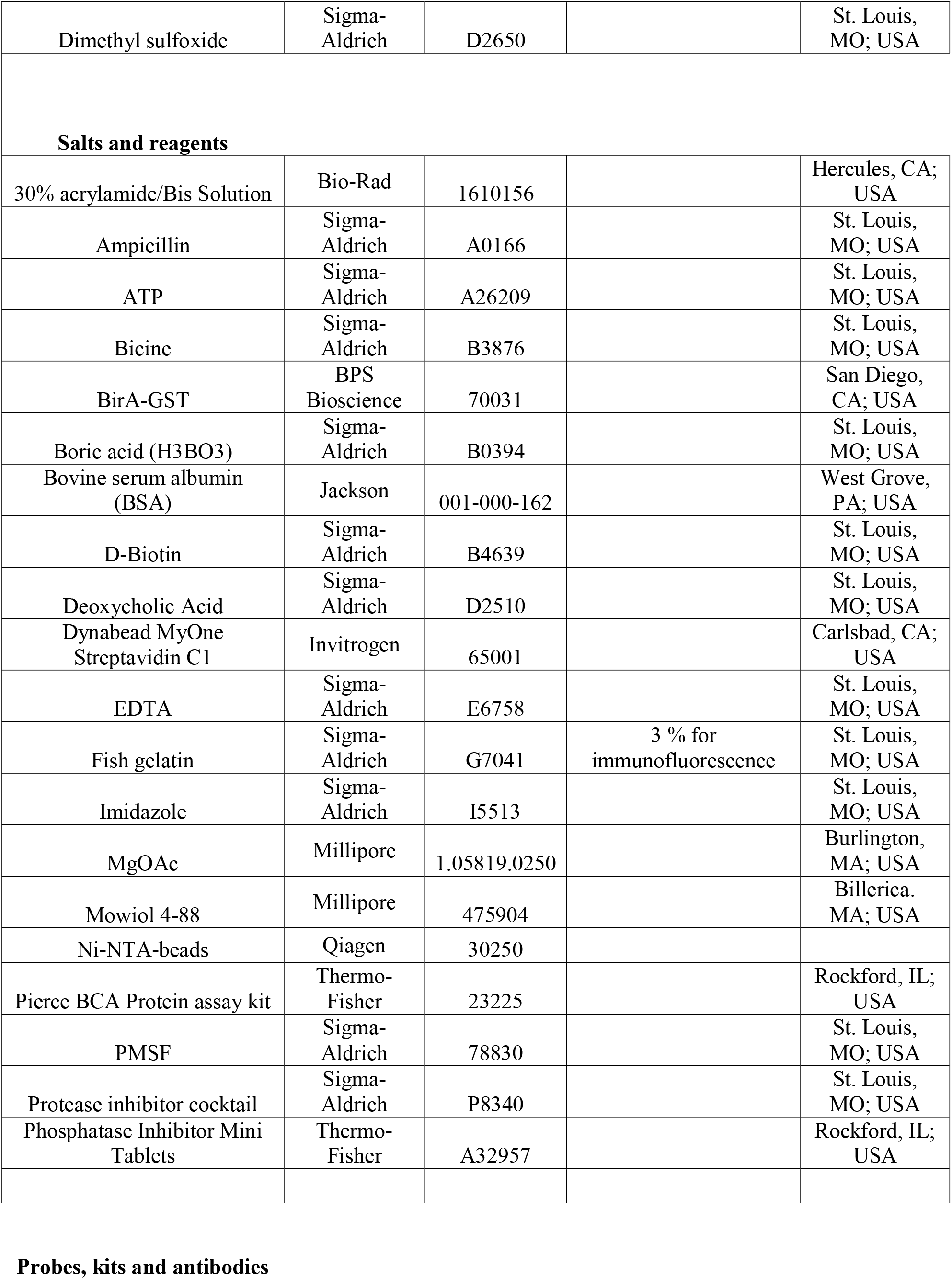
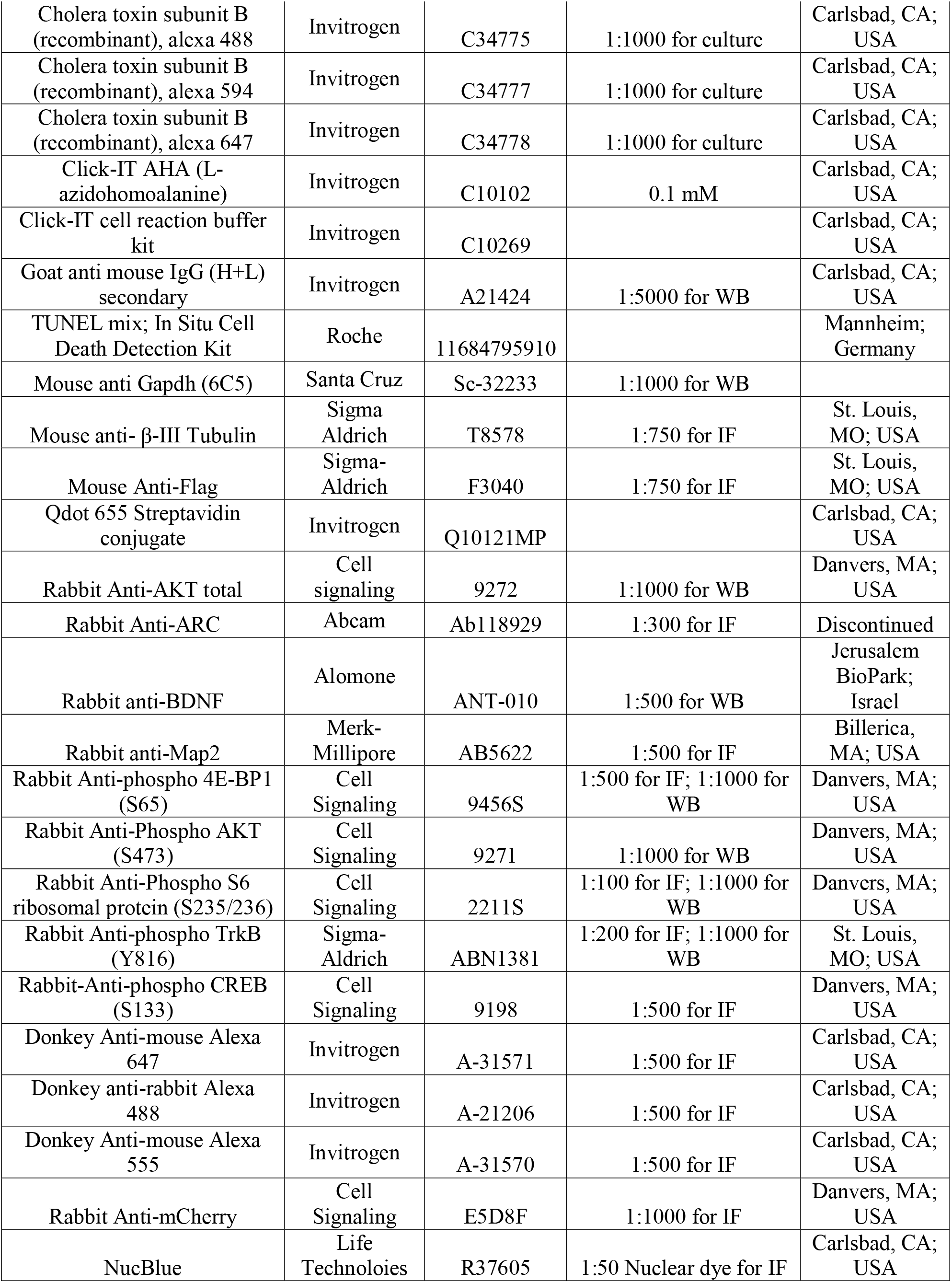
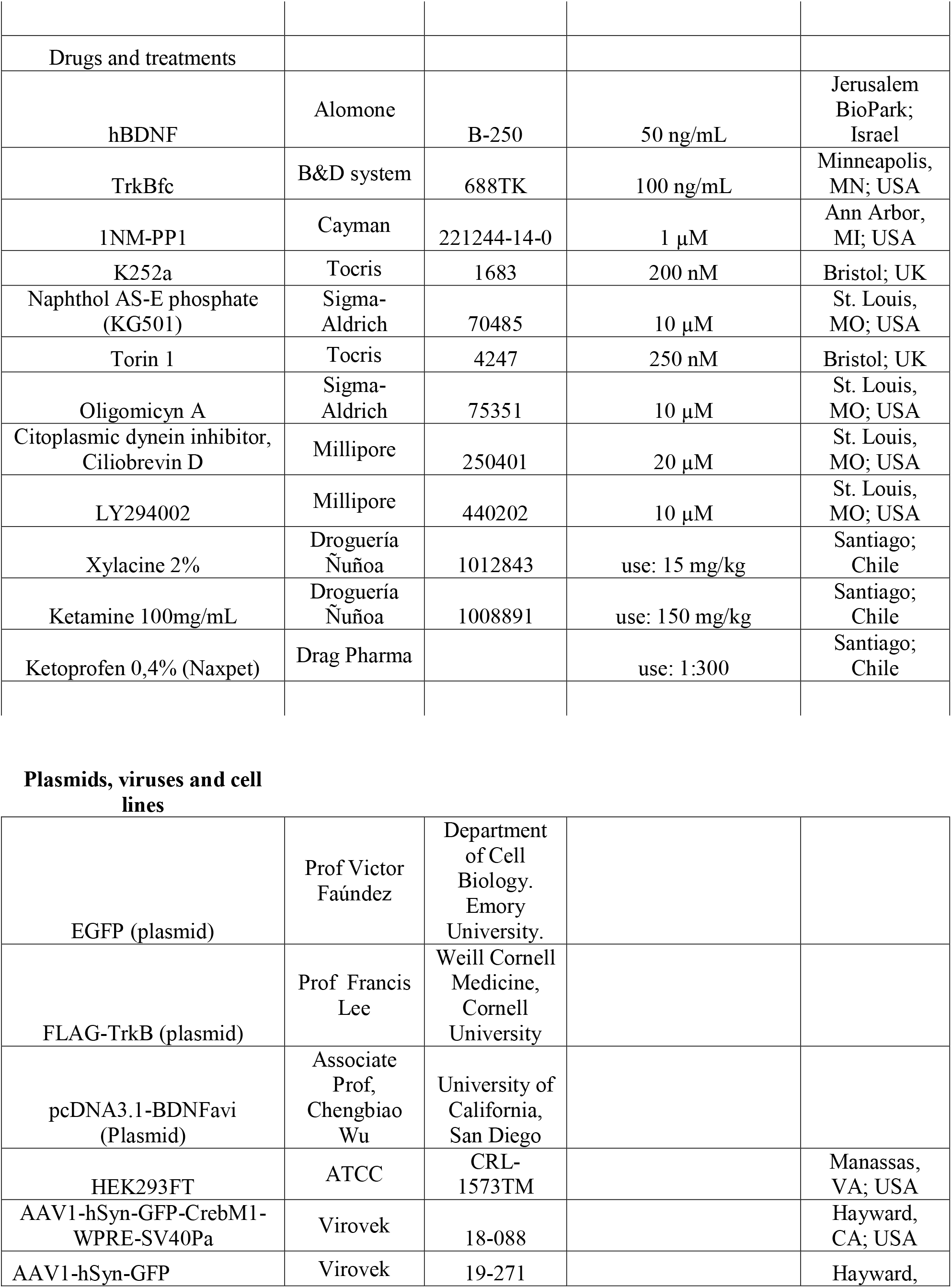
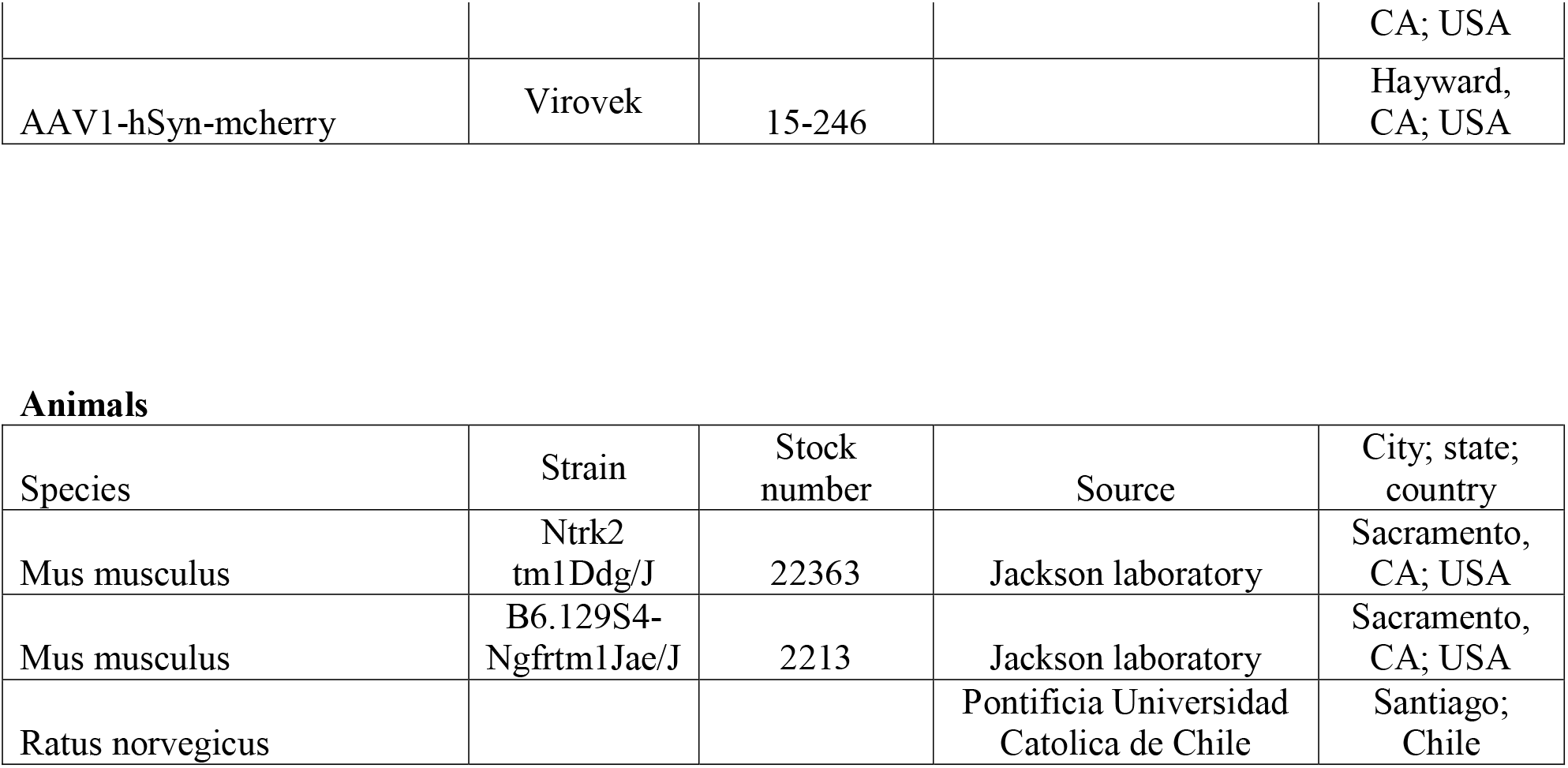
List of materials.

## REFERENCES

1. Andres-Alonso, M., Ammar, M. R., Butnaru, I., Gomes, G. M., Acuna Sanhueza, G., Raman, R., Yuanxiang, P., Borgmeyer, M., Lopez-Rojas, J., Raza, S. A., Brice, N., Hausrat, T. J., Macharadze, T., Diaz-Gonzalez, S., Carlton, M., Failla, A. V., Stork, O., Schweizer, M., Gundelfinger, E. D., Kneussel, M., Spilker, C., Karpova, A. & Kreutz, M. R. 2019. SIPA1L2 controls trafficking and local signaling of TrkB-containing amphisomes at presynaptic terminals. Nat Commun, 10, 5448.

2. Andreska, T., Rauskolb, S., Schukraft, N., Lüningschrör, P., Sasi, M., Signoret-Genest, J., Behringer, M., Blum, R., Sauer, M., Tovote, P. & Sendtner, M. 2020. Induction of BDNF Expression in Layer II/III and Layer V Neurons of the Motor Cortex Is Essential for Motor Learning. J Neurosci, 40, 6289–6308.

3. Barco, A., Patterson, S. L., Alarcon, J. M., Gromova, P., Mata-Roig, M., Morozov, A. & Kandel, E. R. 2005. Gene expression profiling of facilitated L-LTP in VP16-CREB mice reveals that BDNF is critical for the maintenance of LTP and its synaptic capture. Neuron, 48, 123–37.

4. Best, J. L., Amezcua, C. A., Mayr, B., Flechner, L., Murawsky, C. M., Emerson, B., Zor, T., Gardner, K. H. & Montminy, M. 2004. Identification of small-molecule antagonists that inhibit an activator: coactivator interaction. Proc Natl Acad Sci U S A, 101, 17622–7.

5. Bonni, A., Brunet, A., West, A. E., Datta, S. R., Takasu, M. A. & Greenberg, M. E. 1999. Cell survival promoted by the Ras-MAPK signaling pathway by transcription-dependent and -independent mechanisms. Science, 286, 1358–62.

6. Bronfman, F. C., Lazo, O. M., Flores, C. & Escudero, C. A. 2014. Spatiotemporal intracellular dynamics of neurotrophin and its receptors. Implications for neurotrophin signaling and neuronal function. Handb Exp Pharmacol, 220, 33–65.

7. Caracciolo, L., Marosi, M., Mazzitelli, J., Latifi, S., Sano, Y., Galvan, L., Kawaguchi, R., Holley, S., Levine, M. S., Coppola, G., Portera-Cailliau, C., Silva, A. J. & Carmichael, S. T. 2018. CREB controls cortical circuit plasticity and functional recovery after stroke. Nat Commun, 9, 2250.

8. Chen, X., Ye, H., Kuruvilla, R., Ramanan, N., Scangos, K. W., Zhang, C., Johnson, N. M., England, P. M., Shokat, K. M. & Ginty, D. D. 2005. A chemical-genetic approach to studying neurotrophin signaling. Neuron, 46, 13–21.

9. Cheng, X. T., Zhou, B., Lin, M. Y., Cai, Q. & Sheng, Z. H. 2015. Axonal autophagosomes use the ride-on service for retrograde transport toward the soma. Autophagy, 11, 1434–6.

10. Cheung, Z. H., Chin, W. H., Chen, Y., Ng, Y. P. & Ip, N. Y. 2007. Cdk5 Is Involved in BDNF-Stimulated Dendritic Growth in Hippocampal Neurons. PLoS Biol, 5, e63.

11. Cohen, M. S., Bas Orth, C., Kim, H. J., Jeon, N. L. & Jaffrey, S. R. 2011. Neurotrophin-mediated dendrite-to-nucleus signaling revealed by microfluidic compartmentalization of dendrites. Proc Natl Acad Sci U S A, 108, 11246–51.

12. Cosker, K. E. & Segal, R. A. 2014. Neuronal signaling through endocytosis. Cold Spring Harb Perspect Biol, 6.

13. Debaisieux, S., Encheva, V., Chakravarty, P., Snijders, A. P. & Schiavo, G. 2016. Analysis of Signaling Endosome Composition and Dynamics Using SILAC in Embryonic Stem Cell-Derived Neurons. Mol Cell Proteomics, 15, 542–57.

14. Deinhardt, K., Salinas, S., Verastegui, C., Watson, R., Worth, D., Hanrahan, S., Bucci, C. & Schiavo, G. 2006. Rab5 and Rab7 control endocytic sorting along the axonal retrograde transport pathway. Neuron, 52, 293–305.

15. Dijkhuizen, P. A. & Ghosh, A. 2005. BDNF regulates primary dendrite formation in cortical neurons via the PI3-kinase and MAP kinase signaling pathways. J Neurobiol, 62, 278–88.

16. Dolmetsch, R. E., Pajvani, U., Fife, K., Spotts, J. M. & Greenberg, M. E. 2001. Signaling to the nucleus by an L-type calcium channel-calmodulin complex through the MAP kinase pathway. Science, 294, 333–9.

17. Escudero, C. A., Cabeza, C., Moya-Alvarado, G., Maloney, M. T., Flores, C. M., Wu, C., Court, F. A., Mobley, W. C. & Bronfman, F. C. 2019. c-Jun N-terminal kinase (JNK)-dependent internalization and Rab5-dependent endocytic sorting mediate long-distance retrograde neuronal death induced by axonal BDNF-p75 signaling. Sci Rep, 9, 6070.

18. Ferreira, T. A., Blackman, A. V., Oyrer, J., Jayabal, S., Chung, A. J., Watt, A. J., Sjöström, P. J. & Van Meyel, D. J. 2014. Neuronal morphometry directly from bitmap images. Nature Methods, 11, 982–984.

19. Finkbeiner, S., Tavazoie, S. F., Maloratsky, A., Jacobs, K. M., Harris, K. M. & Greenberg, M. E. 1997. CREB: a major mediator of neuronal neurotrophin responses. Neuron, 19, 1031–47.

20. Gonzalez-Gutierrez, A., Lazo, O. M. & Bronfman, F. C. 2020. The Rab5-Rab11 Endosomal Pathway is Required for BDNF-Induced CREB Transcriptional Regulation in Hippocampal Neurons. J Neurosci, 40, 8042–8054.

21. Goto-Silva, L., Mcshane, M. P., Salinas, S., Kalaidzidis, Y., Schiavo, G. & Zerial, M. 2019. Retrograde transport of Akt by a neuronal Rab5-APPL1 endosome. Sci Rep, 9, 2433.

22. Grimes, M. L., Zhou, J., Beattie, E. C., Yuen, E. C., Hall, D. E., Valletta, J. S., Topp, K. S., Lavail, J. H., Bunnett, N. W. & Mobley, W. C. 1996. Endocytosis of activated TrkA: evidence that nerve growth factor induces formation of signaling endosomes. J Neurosci, 16, 7950–64.

23. Harrington, A. W. & Ginty, D. D. 2013. Long-distance retrograde neurotrophic factor signalling in neurons. Nat Rev Neurosci, 14, 177–87.

24. Herzog, J. J., Xu, W., Deshpande, M., Rahman, R., Suib, H., Rodal, A. A., Rosbash, M. & Paradis, S. 2020. TDP-43 dysfunction restricts dendritic complexity by inhibiting CREB activation and altering gene expression. Proc Natl Acad Sci U S A, 117, 11760–11769.

25. Hirokawa, N., Niwa, S. & Tanaka, Y. 2010. Molecular motors in neurons: transport mechanisms and roles in brain function, development, and disease. Neuron, 68, 610–38.

26. Horch, H. W. & Katz, L. C. 2002. BDNF release from single cells elicits local dendritic growth in nearby neurons. Nat Neurosci, 5, 1177–84.

27. Huang, E. J. & Reichardt, L. F. 2003. Trk receptors: roles in neuronal signal transduction. Annu Rev Biochem, 72, 609–42.

28. Ibanez, C. F. & Simi, A. 2012. p75 neurotrophin receptor signaling in nervous system injury and degeneration: paradox and opportunity. Trends Neurosci, 35, 431–40.

29. Jan, Y. N. & Jan, L. Y. 2010. Branching out: mechanisms of dendritic arborization. Nat Rev Neurosci, 11, 316–28.

30. Kraemer, B. R., Yoon, S. O. & Carter, B. D. 2014. The biological functions and signaling mechanisms of the p75 neurotrophin receptor. Handb Exp Pharmacol, 220, 121–64.

31. Kumar, V., Zhang, M. X., Swank, M. W., Kunz, J. & Wu, G. Y. 2005. Regulation of dendritic morphogenesis by Ras-PI3K-Akt-mTOR and Ras-MAPK signaling pathways. J Neurosci, 25, 11288–99.

32. Kuruvilla, R., Ye, H. & Ginty, D. D. 2000. Spatially and functionally distinct roles of the PI3-K effector pathway during NGF signaling in sympathetic neurons. Neuron, 27, 499–512.

33. Kwon, M., Fernandez, J. R., Zegarek, G. F., Lo, S. B. & Firestein, B. L. 2011. BDNF-promoted increases in proximal dendrites occur via CREB-dependent transcriptional regulation of cypin. J Neurosci, 31, 9735–45.

34. Lazo, O. M., Gonzalez, A., Ascano, M., Kuruvilla, R., Couve, A. & Bronfman, F. C. 2013. BDNF regulates Rab11-mediated recycling endosome dynamics to induce dendritic branching. J Neurosci, 33, 6112–22.

35. Leal, G., Comprido, D. & Duarte, C. B. 2014. BDNF-induced local protein synthesis and synaptic plasticity. Neuropharmacology, 76 Pt C, 639–56.

36. Lee, K. F., Li, E., Huber, L. J., Landis, S. C., Sharpe, A. H., Chao, M. V. & Jaenisch, R. 1992. Targeted mutation of the gene encoding the low affinity NGF receptor p75 leads to deficits in the peripheral sensory nervous system. Cell, 69, 737–49.

37. Lehigh, K. M., West, K. M. & Ginty, D. D. 2017. Retrogradely Transported TrkA Endosomes Signal Locally within Dendrites to Maintain Sympathetic Neuron Synapses. Cell Rep, 19, 86–100.

38. Liu, Q., Chang, J. W., Wang, J., Kang, S. A., Thoreen, C. C., Markhard, A., Hur, W., Zhang, J., Sim, T., Sabatini, D. M. & Gray, N. S. 2010. Discovery of 1-(4-(4-propionylpiperazin-1-yl)-3-(trifluoromethyl)phenyl)-9-(quinolin-3-yl)benz o[h][1,6]naphthyridin-2(1H)-one as a highly potent, selective mammalian target of rapamycin (mTOR) inhibitor for the treatment of cancer. J Med Chem, 53, 7146–55.

39. Lom, B., Cogen, J., Sanchez, A. L., Vu, T. & Cohen-Cory, S. 2002. Local and target-derived brain-derived neurotrophic factor exert opposing effects on the dendritic arborization of retinal ganglion cells in vivo. J Neurosci, 22, 7639–49.

40. Lonze, B. E. & Ginty, D. D. 2002. Function and regulation of CREB family transcription factors in the nervous system. Neuron, 35, 605–23.

41. Matsuda, N., Lu, H., Fukata, Y., Noritake, J., Gao, H., Mukherjee, S., Nemoto, T., Fukata, M. & Poo, M. M. 2009. Differential activity-dependent secretion of brain-derived neurotrophic factor from axon and dendrite. J Neurosci, 29, 14185–98.

42. Minichiello, L., Calella, A. M., Medina, D. L., Bonhoeffer, T., Klein, R. & Korte, M. 2002. Mechanism of TrkB-mediated hippocampal long-term potentiation. Neuron, 36, 121–37.

43. Moya-Alvarado, G., Gonzalez, A., Stuardo, N. & Bronfman, F. C. 2018. Brain-Derived Neurotrophic Factor (BDNF) Regulates Rab5-Positive Early Endosomes in Hippocampal Neurons to Induce Dendritic Branching. Front Cell Neurosci, 12, 493.

44. Numrich, J. & Ungermann, C. 2014. Endocytic Rabs in membrane trafficking and signaling. Biological Chemistry, 395, 327–333.

45. Olenick, M. A., Dominguez, R. & Holzbaur, E. L. F. 2019. Dynein activator Hook1 is required for trafficking of BDNF-signaling endosomes in neurons. J Cell Biol, 218, 220–233.

46. Pathak, A., Clark, S., Bronfman, F. C., Deppmann, C. D. & Carter, B. D. 2021. Long-distance regressive signaling in neural development and disease. Wiley Interdiscip Rev Dev Biol, 10, e382.

47. Ravindran, S., Nalavadi, V. C. & Muddashetty, R. S. 2019. BDNF Induced Translation of Limk1 in Developing Neurons Regulates Dendrite Growth by Fine-Tuning Cofilin1 Activity. Front Mol Neurosci, 12, 64.

48. Reck-Peterson, S. L., Redwine, W. B., Vale, R. D. & Carter, A. P. 2018. The cytoplasmic dynein transport machinery and its many cargoes. Nat Rev Mol Cell Biol, 19, 382–398.

49. Redmond, L., Kashani, A. H. & Ghosh, A. 2002. Calcium regulation of dendritic growth via CaM kinase IV and CREB-mediated transcription. Neuron, 34, 999–1010.

50. Riccio, A., Pierchala, B. A., Ciarallo, C. L. & Ginty, D. D. 1997. An NGF-TrkA-mediated retrograde signal to transcription factor CREB in sympathetic neurons. Science, 277, 1097–100.

51. Sainath, R. & Gallo, G. 2015. The dynein inhibitor Ciliobrevin D inhibits the bidirectional transport of organelles along sensory axons and impairs NGF-mediated regulation of growth cones and axon branches. Dev Neurobiol, 75, 757–77.

52. Schindelin, J., Arganda-Carreras, I., Frise, E., Kaynig, V., Longair, M., Pietzsch, T., Preibisch, S., Rueden, C., Saalfeld, S., Schmid, B., Tinevez, J.-Y., White, D. J., Hartenstein, V., Eliceiri, K., Tomancak, P. & Cardona, A. 2012. Fiji: an open-source platform for biological-image analysis. Nature Methods, 9, 676–682.

53. Schratt, G. M., Nigh, E. A., Chen, W. G., Hu, L. & Greenberg, M. E. 2004. BDNF regulates the translation of a select group of mRNAs by a mammalian target of rapamycin-phosphatidylinositol 3-kinase-dependent pathway during neuronal development. J Neurosci, 24, 7366–77.

54. Scott-Solomon, E. & Kuruvilla, R. 2018. Mechanisms of neurotrophin trafficking via Trk receptors. Mol Cell Neurosci, 91, 25–33.

55. Shelton, D. L., Sutherland, J., Gripp, J., Camerato, T., Armanini, M. P., Phillips, H. S., Carroll, K., Spencer, S. D. & Levinson, A. D. 1995. Human trks: molecular cloning, tissue distribution, and expression of extracellular domain immunoadhesins. J Neurosci, 15, 477–91.

56. Shimada, A., Mason, C. A. & Morrison, M. E. 1998. TrkB signaling modulates spine density and morphology independent of dendrite structure in cultured neonatal Purkinje cells. J Neurosci, 18, 8559–70.

57. Sholl, D. A. 1953. Dendritic organization in the neurons of the visual and motor cortices of the cat. J Anat, 87, 387–406.

58. Steward, O. & Worley, P. 2002. Local synthesis of proteins at synaptic sites on dendrites: role in synaptic plasticity and memory consolidation? Neurobiol Learn Mem, 78, 508–27.

59. Stuardo, N., Moya-Alvarado, G., Ramirez, C., Schiavo, G. & Bronfman, F. C. 2020. An Improved Protocol to Purify and Directly Mono-Biotinylate Recombinant BDNF in a Tube for Cellular Trafficking Studies in Neurons. J Vis Exp.

60. Sutton, M. A. & Schuman, E. M. 2006. Dendritic protein synthesis, synaptic plasticity, and memory. Cell, 127, 49–58.

61. Takei, N., Inamura, N., Kawamura, M., Namba, H., Hara, K., Yonezawa, K. & Nawa, H. 2004. Brain-derived neurotrophic factor induces mammalian target of rapamycin-dependent local activation of translation machinery and protein synthesis in neuronal dendrites. J Neurosci, 24, 9760–9.

62. Tapley, P., Lamballe, F. & Barbacid, M. 1992. K252a is a selective inhibitor of the tyrosine protein kinase activity of the trk family of oncogenes and neurotrophin receptors. Oncogene, 7, 371–81.

63. Taylor, A. M., Rhee, S. W., Tu, C. H., Cribbs, D. H., Cotman, C. W. & Jeon, N. L. 2003. Microfluidic Multicompartment Device for Neuroscience Research. Langmuir, 19, 1551–1556.

64. Watakabe, A., Takaji, M., Kato, S., Kobayashi, K., Mizukami, H., Ozawa, K., Ohsawa, S., Matsui, R., Watanabe, D. & Yamamori, T. 2014. Simultaneous visualization of extrinsic and intrinsic axon collaterals in Golgi-like detail for mouse corticothalamic and corticocortical cells: a double viral infection method. Front Neural Circuits, 8, 110.

65. Watson, F. L., Heerssen, H. M., Moheban, D. B., Lin, M. Z., Sauvageot, C. M., Bhattacharyya, A., Pomeroy, S. L. & Segal, R. A. 1999. Rapid nuclear responses to target-derived neurotrophins require retrograde transport of ligand-receptor complex. J Neurosci, 19, 7889–900.

66. Wong, R. O. & Ghosh, A. 2002. Activity-dependent regulation of dendritic growth and patterning. Nat Rev Neurosci, 3, 803–12.

67. Xu, B., Zang, K., Ruff, N. L., Zhang, Y. A., Mcconnell, S. K., Stryker, M. P. & Reichardt, L. F. 2000. Cortical degeneration in the absence of neurotrophin signaling: dendritic retraction and neuronal loss after removal of the receptor TrkB. Neuron, 26, 233–45.

68. Yamashita, N., Joshi, R., Zhang, S., Zhang, Z. Y. & Kuruvilla, R. 2017. Phospho-Regulation of Soma-to-Axon Transcytosis of Neurotrophin Receptors. Dev Cell, 42, 626–639 e5.

69. Yan, Q., Rosenfeld, R. D., Matheson, C. R., Hawkins, N., Lopez, O. T., Bennett, L. & Welcher, A. A. 1997. Expression of brain-derived neurotrophic factor protein in the adult rat central nervous system. Neuroscience, 78, 431–48.

70. Yap, C. C., Digilio, L., Mcmahon, L. P., Garcia, A. D. R. & Winckler, B. 2018. Degradation of dendritic cargos requires Rab7-dependent transport to somatic lysosomes. J Cell Biol, 217, 3141–3159.

71. Yap, E. L. & Greenberg, M. E. 2018. Activity-Regulated Transcription: Bridging the Gap between Neural Activity and Behavior. Neuron, 100, 330–348.

72. Ying, S. W., Futter, M., Rosenblum, K., Webber, M. J., Hunt, S. P., Bliss, T. V. & Bramham, C. R. 2002. Brain-derived neurotrophic factor induces long-term potentiation in intact adult hippocampus: requirement for ERK activation coupled to CREB and upregulation of Arc synthesis. J Neurosci, 22, 1532–40.

73. Zagrebelsky, M., Holz, A., Dechant, G., Barde, Y. A., Bonhoeffer, T. & Korte, M. 2005. The p75 neurotrophin receptor negatively modulates dendrite complexity and spine density in hippocampal neurons. J Neurosci, 25, 9989–99.

74. Zhao, X., Zhou, Y., Weissmiller, A. M., Pearn, M. L., Mobley, W. C. & Wu, C. 2014. Real-time imaging of axonal transport of quantum dot-labeled BDNF in primary neurons. J Vis Exp, 51899.

75. Zheng, J., Shen, W. H., Lu, T. J., Zhou, Y., Chen, Q., Wang, Z., Xiang, T., Zhu, Y. C., Zhang, C., Duan, S. & Xiong, Z. Q. 2008. Clathrin-dependent endocytosis is required for TrkB-dependent Akt-mediated neuronal protection and dendritic growth. J Biol Chem, 283, 13280–8.

76. Zhou, B., Cai, Q., Xie, Y. & Sheng, Z. H. 2012. Snapin recruits dynein to BDNF-TrkB signaling endosomes for retrograde axonal transport and is essential for dendrite growth of cortical neurons. Cell Rep, 2, 42–51.

77. Zhou, P., Porcionatto, M., Pilapil, M., Chen, Y., Choi, Y., Tolias, K. F., Bikoff, J. B., Hong, E. J., Greenberg, M. E. & Segal, R. A. 2007. Polarized signaling endosomes coordinate BDNF-induced chemotaxis of cerebellar precursors. Neuron, 55, 53–68.

